# Using *Cupriavidus necator* H16 to provide a roadmap for increasing electroporation efficiency in non-model bacteria

**DOI:** 10.1101/2024.05.27.596136

**Authors:** Matteo Vajente, Riccardo Clerici, Hendrik Ballerstedt, Lars M. Blank, Sandy Schmidt

## Abstract

Bacteria are a treasure trove of metabolic reactions, but most industrial biotechnology applications rely on a limited set of established host organisms. In contrast, adopting non-model bacteria for the production of various chemicals of interest is often hampered by their limited genetic amenability coupled with their low transformation efficiency. In this study, we propose a series of steps that can be taken to increase electroporation efficiency in non-model bacteria. As a test strain, we use *Cupriavidus necator* H16, a lithoautotrophic bacterium that has been engineered to produce a wide range of products from CO_2_ and hydrogen. However, its low electroporation efficiency hinders the high-throughput genetic modifications required to develop *C. necator* into an industrially relevant host organism. First, we propose a species-independent technique based on natively methylated DNA and Golden Gate assembly to increase one-pot cloning and electroporation efficiency by 70-fold. Second, bioinformatic tools were used to predict defense systems and develop a restriction avoidance strategy that was used to introduce suicide plasmids by electroporation to obtain a domesticated strain. The results are discussed in the context of metabolic engineering of non-model bacteria.

**TABLE OF CONTENT:** 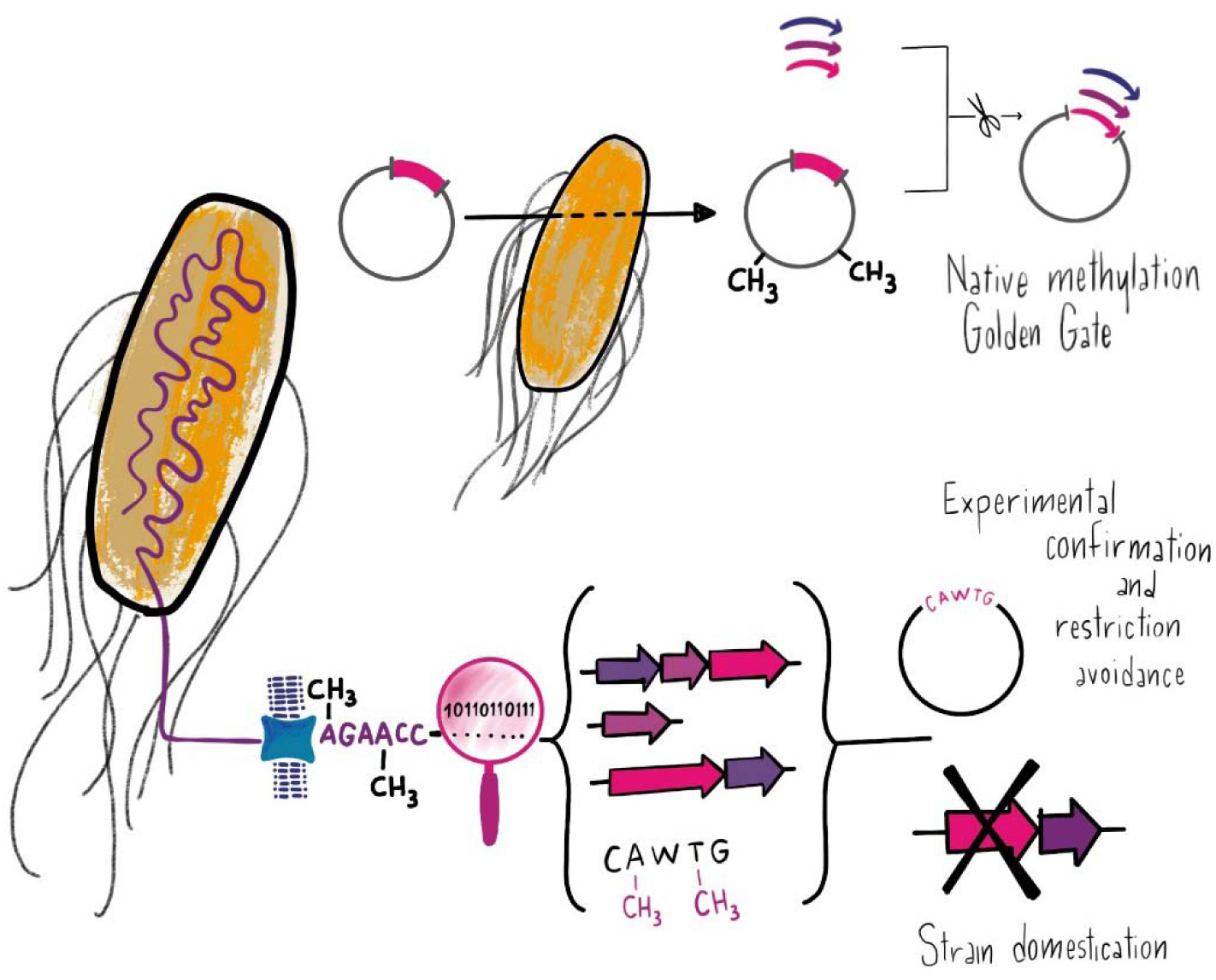

## INTRODUCTION

The concentration of carbon dioxide in the atmosphere is increasing rapidly, and many different actors in the public and private sectors are rushing to develop new technologies to either store it (carbon capture and storage, CCS) or convert it into valuable products (carbon capture and utilization, CCU)^1^. Nature has been operating CCU for a long time, fixing CO_2_ through different metabolic pathways and transforming it into complex organic molecules^2^, making microorganisms a treasure trove of potential CO_2_ devourers^3^. In addition, with the help of synthetic biology tools, these organisms can be modified to produce useful products such as novel foods, fine chemicals, and even bulk chemicals such as 1,3 propanediol^4^. However, most of these tools have been developed for a small selection of domesticated (heterotrophic) bacteria, such as *Escherichia coli*, and less frequently for non-model autotrophic microorganisms.

While *E. coli* or yeasts have been used extensively to produce various chemicals, they cannot natively use CO_2_ as a carbon source. Consequently, there has been much interest in engineering heterotrophic organisms into synthetic autotrophs, but the efficiency and growth rates have been low^5,6^. In contrast, model organisms in the CO_2_-fixing lineages, such as *Synechocystis* (photoautotrophy) or *Cupriavidus necator* (lithoautotrophy), are natively endowed with the CO_2_-fixing machinery. However, their rapid and straightforward genetic engineering remains the bottleneck and often requires time-consuming and extensive resources to integrate pathway optimization, strain optimization and process development^7^.

*Cupriavidus necator* H16 (formerly known as *Ralstonia eutropha* H16) is the most studied hydrogen-oxidizing bacterium, capable of using molecular hydrogen to fuel its aerobic metabolism and fix CO_2_. It has been extensively engineered in the past, and many products have been obtained via metabolic engineering and heterologous enzyme expression^8,9^. Since hydrogen can be obtained via electrolysis using green electricity, this microbe could be used in carbon-negative processes. Interestingly, *C. necator* H16 can also grow on formate as sole carbon and energy source. Formic acid can be obtained from CO_2_ via electrocatalysis and is miscible in water, which overcomes limitations of gas fermentation, such as low solubility of hydrogen^10^. However, there is still a bottleneck in the widespread use of *C. necator* and the application of modern synthetic biology tools. While many plasmids, promoters and ribosome binding sites (RBSs) have been characterized, DNA delivery is mostly based on conjugation from *E. coli* S17-1^11^, which impairs high-throughput genetic modifications, making the engineering process cumbersome and time-consuming. Several research groups have attempted to improve electroporation efficiency or chemical transformation of *C. necator* through protocol optimization^12–14^, plasmid design^12,15^, and strain engineering^16^. However, many recent studies still rely on conjugation for DNA delivery. Thus, DNA delivery remains a critical bottleneck for the application of modern synthetic biology tools. Recent studies suggest that genetic tools and regulatory elements can be exchanged between organisms^17^, and novel strategies have been developed to rapidly identify the plasmid replicons compatible with novel organisms^18^. However, all techniques that rely on DNA expression or modification also require DNA delivery, which often involves laborious efforts to improve conjugation or electroporation protocols. Furthermore, DNA delivery bottlenecks can vary widely even in closely related organisms due to frequent horizontal gene transfer of defense systems^19^.

The main cause of the low electroporation efficiency in *C. necator* is thought to be the presence of various restriction-modification systems (RM systems), which are common in non-domesticated bacteria^16^. In their diverse ecological niches, bacteria use modification enzymes to mark their DNA by methylating nucleotides in specific patterns. This allows cells to distinguish between self and non-self-DNA to avoid phages and other mobile genetic elements (MGEs)^20^. Scientists have devised several strategies to overcome the barrier posed by RM systems: (a) plasmid modification for restriction avoidance: DNA molecules were pre-treated to be recognized as “self” by the host RM system, achieved through *in vitro* methylation^21^ or by using a shuttle strain that correctly modified the DNA^22^; (b) plasmid design for restriction avoidance: by rational elimination of recognition patterns, plasmids could be rendered ‘invisible’ to the bacterial immune system^23^; (c) temporary inactivation of restriction enzymes: in some species, heat-shock pre-treatment temporarily inactivated plasmid restriction^24,25^; (d) deletion of restriction enzymes: cells lacking restriction enzymes have been selected through random mutagenesis or rational engineering, ‘domesticating’ the strain for easier use^26^. All these methods require information about the organism’s genome and methylome, and recently novel algorithms for *de novo* pattern identification have been developed to aid the process^27,28^.

While these examples focused on RM systems, recent advances in the field have led to the discovery of dozens of new defense systems capable of blocking incoming genetic elements. While most of these have only been tested against phages, some operons can also interfere with plasmid transformation or reduce plasmid stability. Examples include the Wadjet system^29^, prokaryotic Argonaute proteins^30^, Cas proteins^31^, and other systems^32^. Bioinformatic tools have also been developed to identify these systems from genomic information^33,34^.

In this study, using *C. necator* H16 as an example, we provide a roadmap (Figure 1) for increasing electroporation efficiency in non-model bacteria by applying different methods and tools to characterize its defense arsenal and ultimately increase electroporation efficiency. First, we developed a new method that allows one-pot cloning and electroporation of natively methylated DNA. This method is technically species-independent and could improve initial screenings in other non-model bacteria. Second, we investigated defense systems in *C. necator* H16 by using bioinformatic tools and databases, and used ad-hoc plasmids to investigate the role of each restriction enzyme during electroporation. Finally, we deleted three promising operons and obtained *C. necator* ΔRM, a strain with increased electroporation efficiency. Thus, we aim to contribute to the improved engineerability of *C. necator* H16 and provide a roadmap for DNA delivery in newly isolated bacteria.

**Figure 1:**
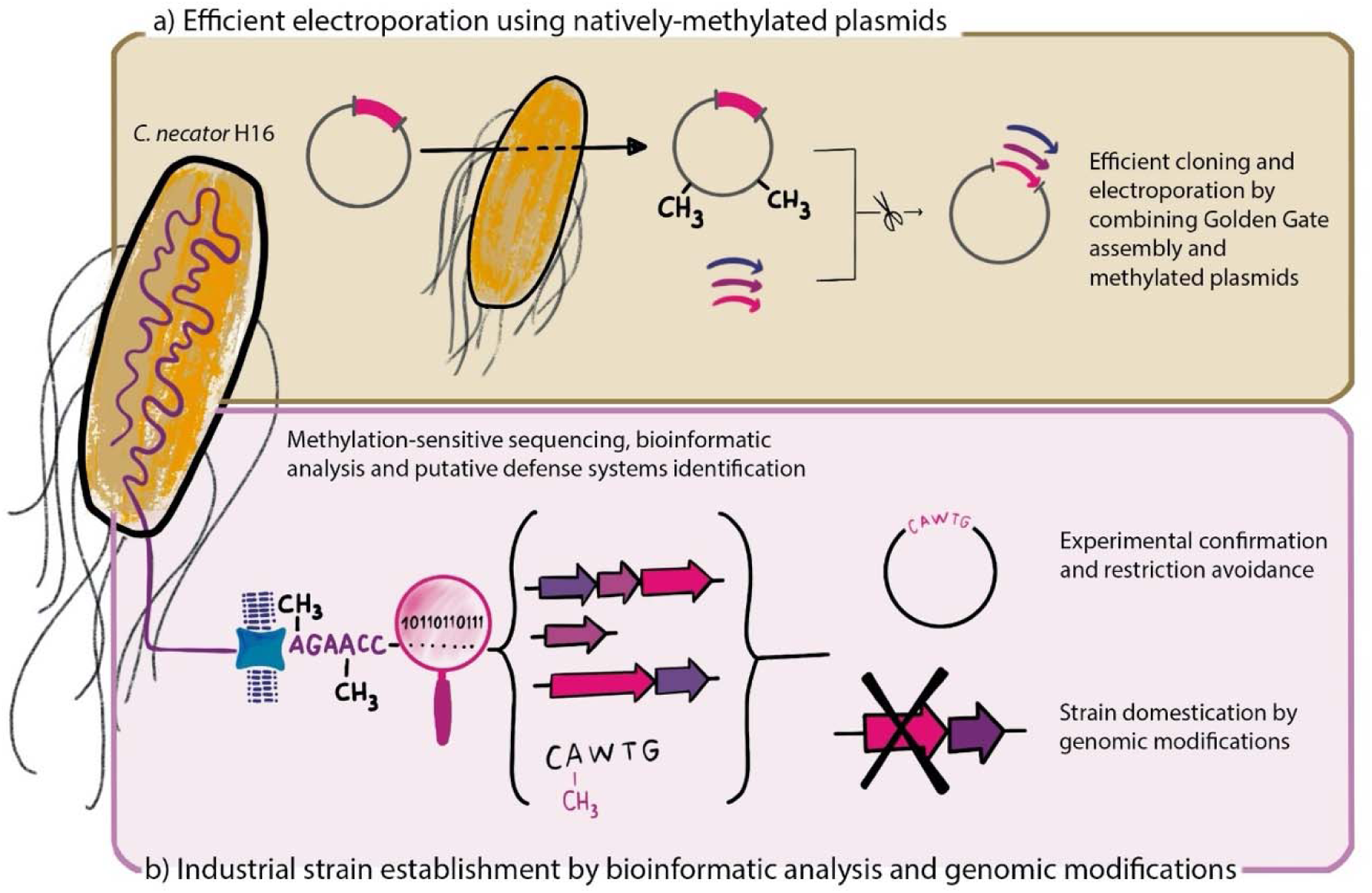
Steps needed to increase the electroporation efficiency in the non-model bacterium *C. necator* H16. A) Using Golden Gate assembly with natively methylated DNA increases electroporation efficiency in *C. necator* H16 without knowing any information on its defense arsenal; b) Analysis and characterization of endogenous defense systems by bioinformatic prediction and experimental confirmation. These systems can then be overcome by restriction avoidance or by rational strain engineering.

## RESULTS

### Restriction-modification systems impair large plasmid electroporation

We first investigated three different electroporation protocols using the small plasmid pCAT201 (3198 bp) as a benchmark (Figure 2A, Table 1)^12,14,15^. We found the protocol derived from Tee et al. (2017) to be the most efficient and used it for all subsequent experiments. During our experiments, we also discovered that plating on LB agar supplemented with 200 mg/L kanamycin sometimes led to the growth of colonies without plasmid (escaper colonies) when plating at high cell density. The issue disappeared when we supplemented 400 mg/L kanamycin, while the colony count did not change (Supplementary Information, Figure S1). From then on, we used the higher concentration for selection after electroporation. We then increased the size of the plasmid and created plasmid pCAT_par (5448 bp) by cloning the partitioning region of plasmid RP4 (*parCBA, parDE*) into pCAT201^35^. Since *C. necator* H16 is known for its plasmid instability, expression plasmids often contain this or other stabilizing regions to ensure correct plasmid segregation^36–39^. The partitioning region contains a toxin-antitoxin operon (*parDE*) and a system that includes a site-specific recombination system mediating resolution of plasmid multimers (*parCBA*)^40,41^.

**Figure 2:**
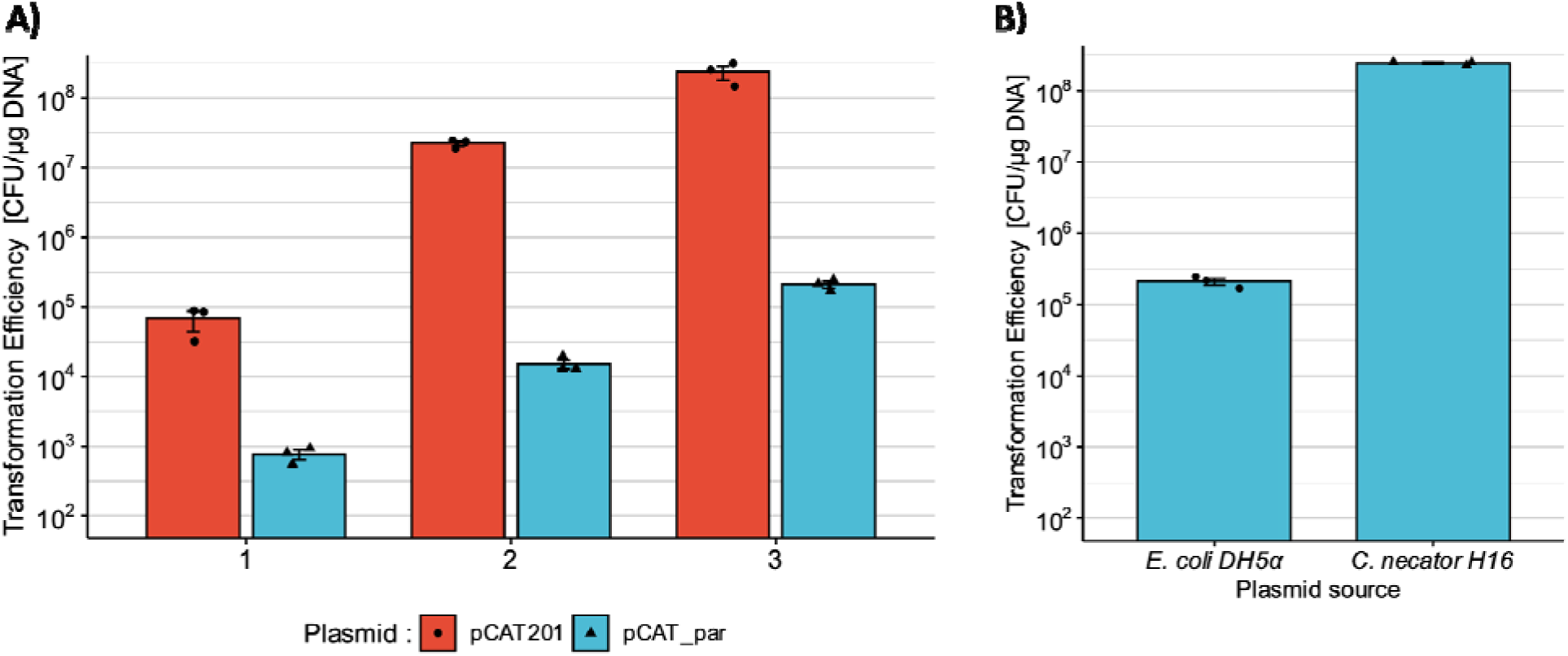
Restriction-Modification systems impair large plasmid electroporation. A) Comparison of three different electroporation protocols using two benchmark plasmids, pCAT201 (red) and pCAT_par (blue). 50 µL of competent *C. necator* H16 cells were transformed with 50 ng of plasmid. The protocols 1, 2 and 3 were derived from Ehsaan et al. (2021)^12^, Azubuike et al. (2021)^15^, and Tee et al. (2017)^14^, respectively. The protocols were modified as described in Materials and Methods and Supplementary Information. B) Transformation efficiencies of the same plasmid (pCAT_par) extracted from *E. coli* DH5a and *C. necator* H16. For each condition tested, three transformations were performed (mean and standard deviation reported).

**Table 1:** Strains and plasmids used in this study.

| Strain | Genotype* | Source or reference |
| --- | --- | --- |
| *C. necator* **H16** /<br>*C. necator* *wt* | *Cupriavidus necator* H16 (DSM 428) | DSMZ |
| *C. necator* **ΔRM** | *C. necator* H16 ΔE6A55_RS00030 ΔE6A55_RS00035<br>ΔE6A55_RS00040 ΔE6A55_RS00045 | This study |
| *C. necator* **Δwad** | *C. necator* H16 ΔE6A55_RS00090 ΔE6A55_RS00095<br>ΔE6A55_RS00100 ΔE6A55_RS00105 | This study |
| *C. necator* **ΔΔ** | *C. necator* Δwad ΔE6A55_RS00030 ΔE6A55_RS00035<br>ΔE6A55_RS00040 ΔE6A55_RS00045 | This study |
| *Escherichia coli* **DH5α** | *F*- Φ80*lacZ*Δ*M15* Δ(*lacZYA*-*argF*) *U169* *recA1* *endA1*<br>*hsdR17*(*rk*-, *mk*+) *phoA* *supE44* *thi-1* *gyrA96* *relA1* λ- | ThermoFisher<br>Scientific |
| *Escherichia coli* **dam-**<br>*/dcm-* | *ara-14* *leuB6* *fhuA31* *lacY1* *tsx78* *glnV44* *galK2* *galT22* *mcrA*<br>*dcm-6* *hisG4* *rfbD1* *R(zgb210::Tn10)* *TetS* *endA1* *rspL136*<br>(*StrR*) *dam13::Tn9* (*CamR*) *xylA-5* *mtl-1* *thi-1* *mcrB1* *hsdR2* | New England Biolabs |
| Plasmid | Characteristics* | Source or reference |
| **pCAT201** | pBBR1 *ori*; *Km<sup>r</sup>*; *lacZ* | Azubuike et al., 2021<br>15 |
| **pCAT_par** | pCAT201; RP4 *parABCDE* | This study |
| **pCATMt** | pCAT201; insertion of 12 nt (GCAAGCAGGCATC) | This study |
| **pMVRha** | pCAT_par; *pJ5[C2]-spisPink*; *rhaRS*; *RP4 oriT* | This study |
| **RSF1010-GFP** | RSF1010 *ori*; *Km<sup>r</sup>*; *J23100-B0034m-GFPmut3* | This study |
| **pLO3** | *Km<sup>r</sup>* *sacB*, RP4 *oniT*, *ColEI ori* | Lenz and Friedrich,<br>1998<sup>42</sup> |
| **pLO3RM** | pLO3; silent mutation in restriction enzyme recognition site; 1<br>kb homology regions | This study |
| **pLO3wad** | pLO3; silent mutation in restriction enzyme recognition site; 1<br>kb homology regions | This study |

\* Detailed strain and plasmid construction are described in the Supplementary Information.

However, while the increase in size was only modest (2250 bp), the loss of electroporation efficiency was significant (Figure 2A). For all protocols, a 100 to 1000-fold loss of electroporation efficiency compared to pCAT201 was observed. While protocol optimization may indeed affect efficiency, these experiments also highlight that the bottleneck for larger plasmids lies elsewhere and we hypothesized a major role for RM systems in this phenomenon. Thus, we compared the natively methylated pCAT_par (extracted from *C. necator*) with an incorrectly methylated one (extracted from *E. coli* DH5α) to confirm that the RM system was the cause of the reduced efficiency (Figure 2B). Indeed, when the plasmid was extracted from the host, the loss of efficiency was completely recovered, and electroporation efficiencies comparable to pCAT201 were achieved. Similar results were briefly mentioned in a previous study^43^. This result suggests a strong involvement of RM systems in electroporation, as we observed that using correctly methylated plasmids drastically improves transformation efficiency. While protocol optimization can improve the electroporation of small plasmids, restriction avoidance is necessary as plasmid size increases.

### The combination of Golden Gate Assembly and natively methylated DNA enables high-efficiency transformation of non-model bacteria

The sharp difference in electroporation efficiency between *E. coli*-methylated and natively methylated DNA has been observed in many non-model bacteria, such as *P. putida*, *P. aeruginosa*^44^, *Actinomyces viscosus*, *Actinomyces naeslundii*^45^, *Caldimonas manganoxidans*^46^, *Campylobacter jejuni*^47^, *Salmonella enteritidis*^48^, and *Staphylococcus carnosus*^25^. Therefore, it can be assumed that RM systems are often the main bottleneck for DNA delivery in bacteria and that they can be avoided by transforming a natively methylated plasmid. We hypothesized that using a plasmid extracted directly from *C. necator* for subsequent cloning would significantly improve electroporation efficiency. Skipping *E. coli* amplification after DNA assembly preserves the host methylation pattern and avoids restriction enzymes. The backbone (extracted from the native host) would be correctly methylated, while the insert would be either unmethylated (PCR amplified) or methylated in the wrong pattern (*E. coli*) (Figure 3A). In this case, the restriction enzymes would only recognize and target the short insert. We decided to further investigate this hypothesis by using Golden Gate assembly, which does not rely on PCR amplification of the backbone (in this case removing the methylation patterns) and can ligate multiple inserts with high efficiency^49^ (Figure 3A).

**Figure 3:**
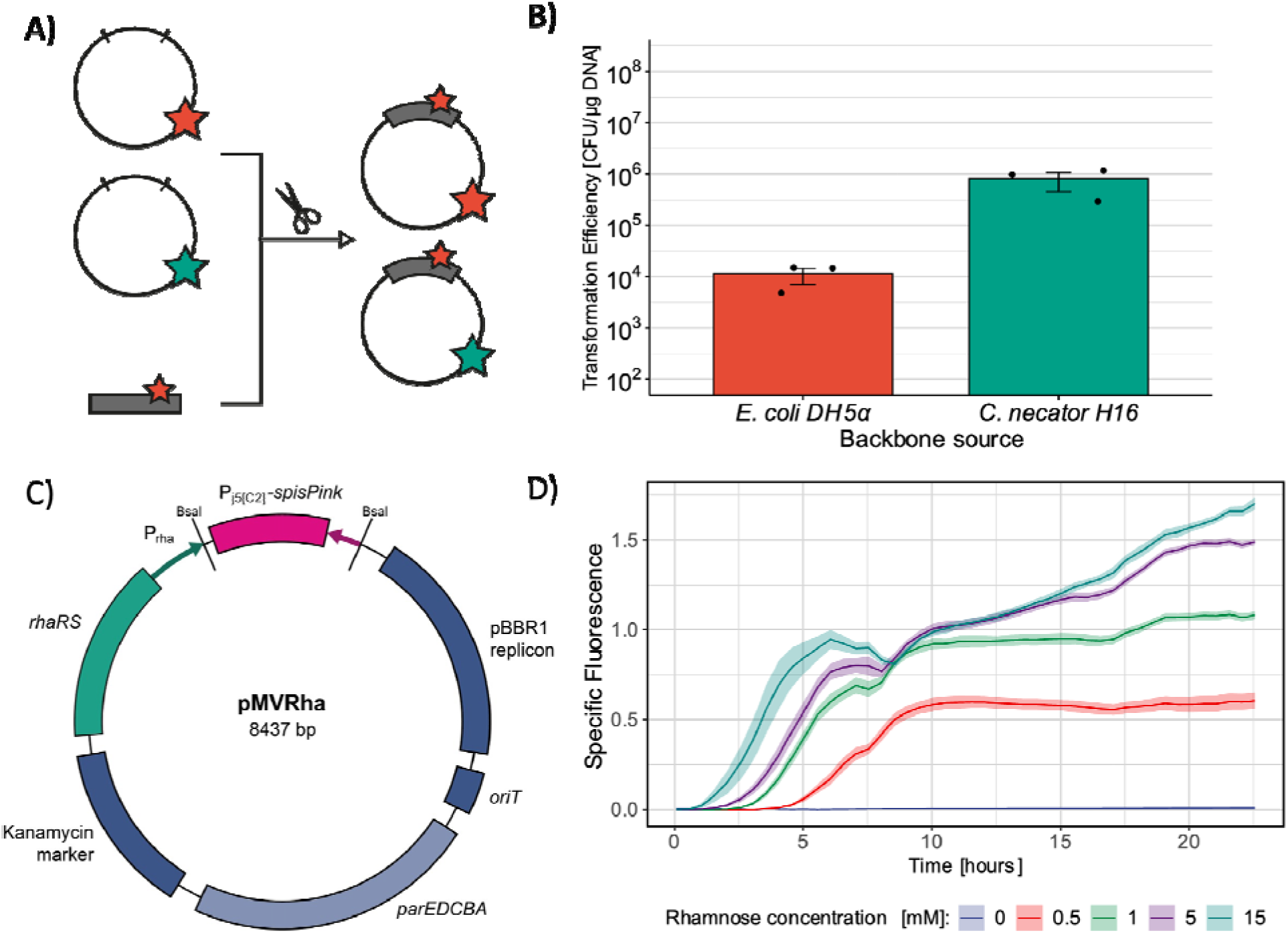
Using Golden Gate assembly with natively methylated DNA increases electroporation efficiency. A) Graphical explanation of the protocol. Plasmid pMVRha is either extracted from *C. necator* H16 (green star) or *E. coli* DH5α (red star). Using a plasmid extracted from the host organism as a backbone creates a final plasmid with a hybrid methylation pattern; B) Efficiency of one-pot Golden Gate assembly and electroporation using as backbone pMVRha extracted from *E. coli* DH5a and *C. necator* H16, respectively. For each condition tested, three Golden Gate reactions and transformations were performed (mean and standard deviation reported); C) Map of the plasmid pMVRha; D) Specific fluorescence of *C. necator* pMVRha-GFP after induction with different concentrations of rhamnose. Five random colonies were picked and tested (mean and standard deviation reported).

First, we created the plasmid pMVRha (8437 bp) (Figure 3C), a Golden Gate-compatible plasmid based on the pCAT201 backbone (Table 1). It contains the *par* stabilizing region^35^, the origin of transfer from RP4 (*oriT*)^11^, a rhamnose-inducible promoter (RhaRS + P_Rha_)^50^, BsaI restriction sites, and a chromoprotein gene as cloning marker (*spisPink*)^51^. Colonies containing un-restricted pMVRha show pink color on plate. Notably, the BsaI overhangs are compatible with all MoClo libraries for easy cloning of genes already available in a donor plasmid^52^.

We then assessed whether this backbone could be used for one-pot Golden Gate assembly and electroporation. We transformed *C. necator* H16 with pMVRha via electroporation, extracted it, and obtained natively methylated DNA. The same plasmid was also extracted from *E. coli* DH5α. We then used these two plasmids as backbones for Golden Gate assembly and cloned the gene *gfpmut3* from a donor plasmid using BsaI^52^. The donor plasmid containing the gene was extracted from *E. coli* DH5α, and 2 µL of each assembly mix were used directly for *C. necator* electroporation. Using the plasmid extracted from *C. necator* as the backbone increased the efficiency 70-fold (Figure 3B), and more than 12000 colonies were obtained after a single electroporation. All colonies were white, confirming successful *spisPink* excision. Thus, using a natively methylated backbone increases efficiency, even if the insert is incorrectly methylated. To confirm that all colonies contained the correct insert, 90 randomly selected colonies showed GFP production after rhamnose induction (Figure S4). Five colonies were randomly selected to confirm the induction parameters, and GFP expression was measured after induction with different concentrations of rhamnose (Figure 3D).

This simple and rapid technique could dramatically improve engineering efforts in non-model bacteria. All organisms compatible with the pBBR1 origin of replication could be transformed with pMVRha by electroporation or conjugation. After this initial transformation, the natively methylated plasmid can be used as a backbone for Golden Gate assembly, ensuring higher electroporation efficiency.

### Bioinformatic tools identify RM systems, restriction sites and other potential transformation bottlenecks

After successfully confirming that RM systems impair plasmid electroporation in *C. necator* H16, we aimed to identify the respective restriction enzymes and their corresponding recognition sequences to develop appropriate strategies to overcome their negative effects on electroporation (e.g., plasmid design, use of pre-methylated plasmid, or temporary restriction inactivation). Knowledge of the genome sequence and the methylome is required to identify the responsible RM systems. Interestingly, both datasets for *C. necator* H16 were already available online at NCBI^53,54^ and REBASE, respectively (Little et al, 2019, REBASE Ref No. 28544).

We analyzed the genome of *C. necator* H16 using three online defense system identification tools, REBASE, DefenseFinder and PADLOC^33,34^. REBASE contains information regarding bacterial methylation and RM systems^53^, while DefenseFinder and PADLOC are both able to predict a vast array of defense systems, including RM enzymes. While some defense systems were identified by only one of the tools, there was significant overlap, allowing us to predict several operons and their potential role in defense (Table S1). We focused our analysis on four of the systems identified (Table 2).

**Table 2:**
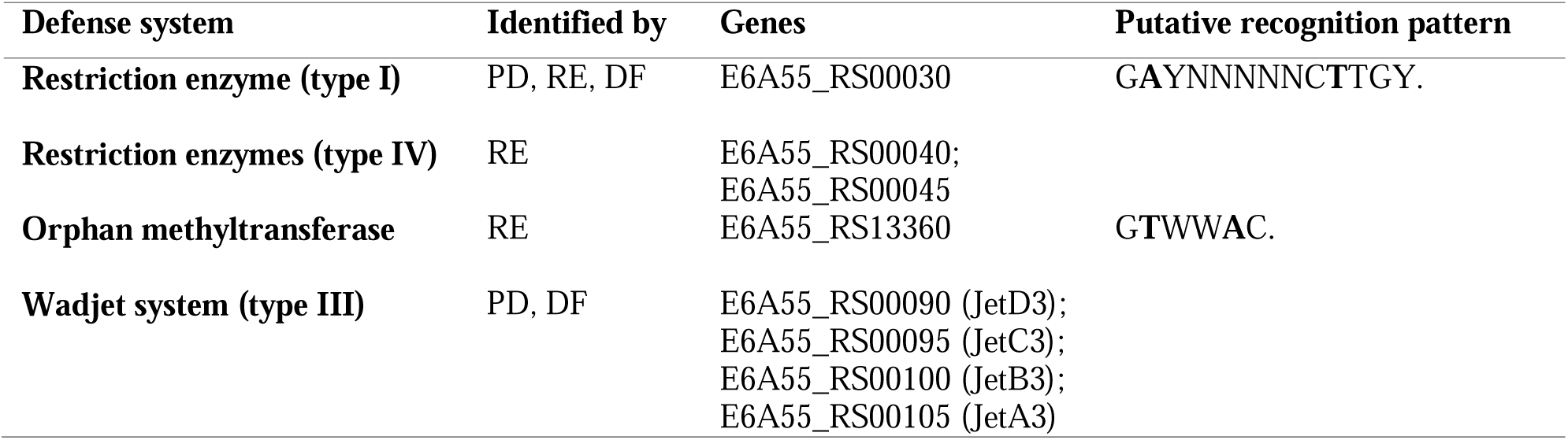
Selected defense systems identified in *C. necator* H16. In “Identified by”: DF: DefenseFinder, RE: REBASE, PD: PADLOC. The putative methylated residues are bold. The complete list can be found in the Supplementary Information (Table S1).

Two methylation patterns were annotated in REBASE, the pattern GAYNNNNNCTTGY is predicted to be methylated (m6A) by a putative type I RM system, which was identified by all algorithms. The pattern GTWWAC is predicted to be methylated by an orphan methylase without an associated restriction enzyme. This lack of restriction element could be caused by incorrect prediction, incomplete horizontal transfer, mutation, or involvement of the methyltransferase in processes unrelated to defense. Two other potential RM enzymes were identified, a type IV restriction enzyme (capable of degrading incorrectly methylated DNA^55^) and another putative orphan methylase (Table S1).

Many other defense systems were identified in the genome, such as AVAST, Zorya, and Gabija (Table S1)^29,56^. However, our attention was focused on the putative Wadjet system, one of the few defense systems that has been shown to act against plasmids by reducing electroporation efficiency and/or decreasing plasmid stability both in their native hosts and when expressed heterologously. When reconstituted *in vitro*, Wadjet systems were able to selectively cut plasmid DNA^29,57–59^. Since *C. necator* H16 is known for its plasmid instability, we hypothesized a role of the putative Wadjet system in plasmid loss.

### Ad-hoc test plasmids to analyze the role of each RM system

To experimentally confirm the bioinformatic predictions of the three tools and clarify the role of these genes in electroporation efficiency, several test plasmids were designed for transformation of *C. necator* H16. First, we created a plasmid to confirm the presence of the type I RM system. We slightly modified pCAT201 by introducing a single restriction pattern (GAYNNNNNCTTGY, 12 bp) and obtained the plasmid pCATMt. When we compared the electroporation efficiency, we observed a drastic loss (5225-fold) with plasmid pCATMt compared to pCAT201 (Figure 4A), although the two plasmids differ by only 12 bp. Therefore, not only is a type I restriction enzyme present, but the pattern predicted in REBASE was also correct.

**Figure 4:**
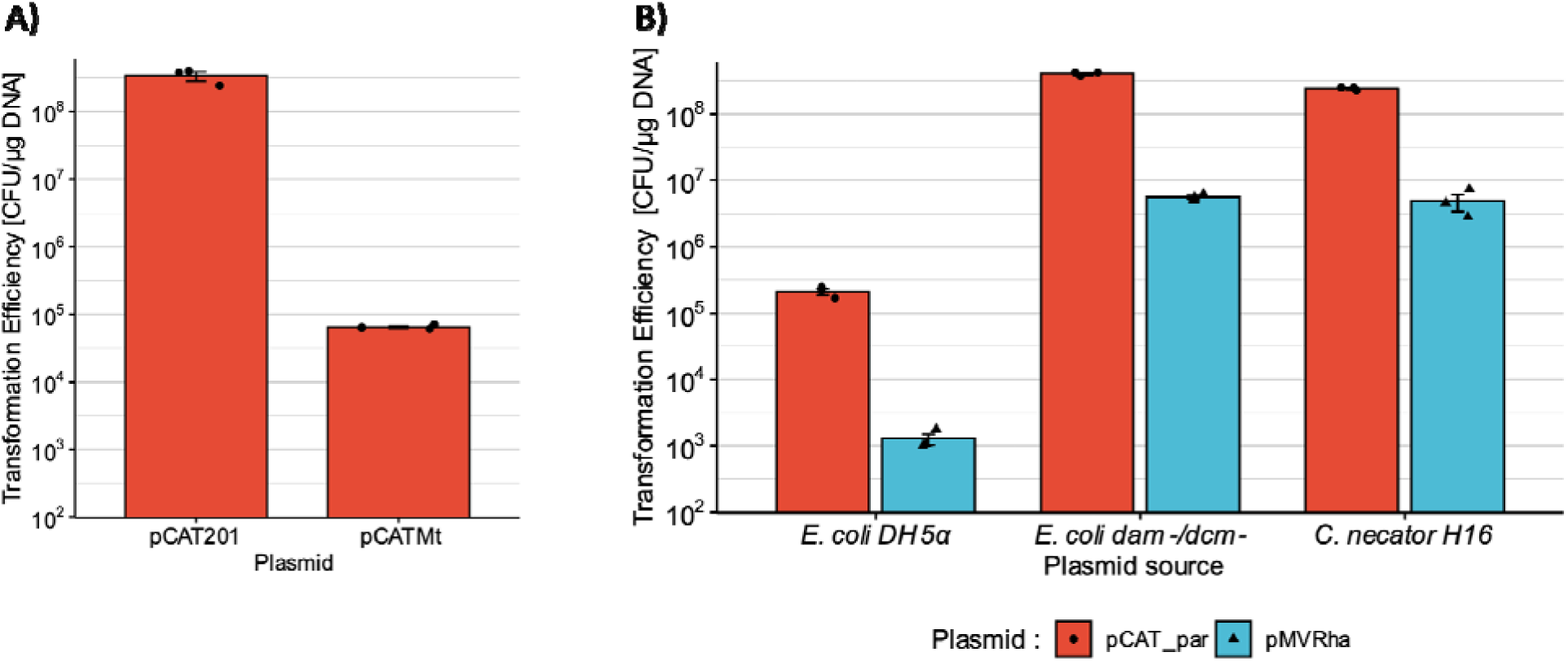
The predicted type I and type IV RM systems are present and decrease electroporation efficiency. A) Transformation efficiency of plasmids pCAT201 and pCATMt in *C. necator* H16; B) Transformation efficiency of two benchmark plasmids (red: pCAT_par; blue: pMVRha) extracted from *E*. *coli* DH5a, *E. coli* dam^-^/dcm^-^, and *C. necator* H16. For each condition tested, three transformations were performed (mean and standard deviation reported).

Interestingly, deletion of the same gene (*H16_A0006*) was the most successful deletion in previous efforts to create a domesticated *C. necator* strain^16^. The transformation efficiency of their test plasmid pBBR1-rfp increased 1658-fold in the deleted strain. However, another group recreated the same strain and obtained more limited results (3-fold increase)^12^. We found that the test plasmid pBBR1-rfp contained the restriction pattern identified by our bioinformatic analysis. This confirms the validity of their observation and explains the results obtained later. The recognition pattern is long and should be rare in plasmids. Notably, many of the plasmids used in *C. necator* H16 contain this pattern (e. g. pKRC, pKRha, pBBR1MCS2)^50,60,61^, which highlights the importance of using different test plasmids to avoid artifacts.

To verify the presence of the type IV restriction system, we used the previously assembled plasmid pCAT_par. This plasmid was chosen for several reasons. First, pCAT_par does not contain the previously identified restriction pattern. Therefore, the contribution of the type I RM system is expected to be neglectable. Second, pCAT_par showed different transformation efficiencies when extracted from *E. coli* and *C. necator*, indicating that another restriction enzyme should be involved. Finally, this plasmid contains a GTWWAC pattern that is methylated in *C. necator*. This pattern was predicted to be methylated by an orphan methylase and may be involved in defense or other mechanisms.

To isolate the role of the type IV restriction enzyme, we transformed pCAT_par in the shuttle strain *E. coli dam^-^/dcm^-^* to obtain unmethylated DNA^62^. We then compared the transformation efficiency of plasmid DNA extracted from *E. coli* DH5α (methylated in the patterns GATC and CCWGG), *E. coli dam^-^/dcm^-^* (not methylated), and *C. necator* (methylated in the patterns GAYNNNNNCTTGY and GTWWAC) (Figure 4B). Interestingly, using unmethylated DNA completely restored transformation efficiency, which was also confirmed with the longer plasmid pMVRha (Figure 4B). Therefore, we assume that only the type I RM system and the type IV system are involved in the loss of electroporation efficiency, and the orphan methylase is involved in defense-unrelated mechanisms. Notably, there is still a decrease in efficiency from pCAT_par to pMVRha even when unmethylated or natively methylated DNA is used.

### Deletion of defense systems by electroporation of suicide plasmids

After confirming the presence of two active RM systems in *C. necator* H16, we created a domesticated strain by gene deletion using the suicide plasmid pLO3, previously used in *C. necator* genome engineering^42^. Suicide plasmids have a lower transformation efficiency than normal plasmids due to the low frequency of genomic integration. We used the previously collected information to transform this plasmid by electroporation. To do so, we used a combination of plasmid engineering and pre-methylation. First, we identified a restriction sequence (GAYNNNNNCTTGY) in the suicide plasmid and used targeted mutagenesis to remove it by introducing a silent mutation. Second, we transformed the shuttle strain *E. coli dam^-^/dcm^-^* with this plasmid to remove DNA methylation. We then successfully transformed *C. necator* by electroporation, although with the expected low efficiency (∼8 CFU/µg DNA). After *sacB*-mediated counter selection and confirmation, we obtained the strain *C. necator* ΔRM (Table 1), where both the putative type I and type IV restriction enzymes were deleted. We used the same technique to delete the Wadjet operon (*C. necator* Δwad) and created a double-deleted strain called *C. necator* Δ (Table 1). Deletions were confirmed via gRNA extraction, PCR amplification of the genomic region and sequencing of the amplicons (Figure S2, S3).

We then characterized these new strains using plasmids pCAT201, pCATMt, pCAT_par, and pMVRha. We performed transformations with plasmids extracted from *E. coli* DH5a and *E. coli dam^-^/dcm^-^*to quantify the role of the two RM systems in each strain (Figure 5). The transformation efficiency of plasmid pCATMt was entirely restored in the *C. necator* ΔRM and ΔΔ knock-out strains. This proved that we had successfully deleted the type I restriction enzyme. Knocking out the type IV restriction enzyme slightly increased the efficiency of the mis-methylated pCAT_par and pMVRha (Figure 5A) but did not completely restore it. In fact, plasmids extracted from *E. coli dam^-^/dcm^-^* achieved higher transformation efficiencies (Figure 5B).

**Figure 5:**
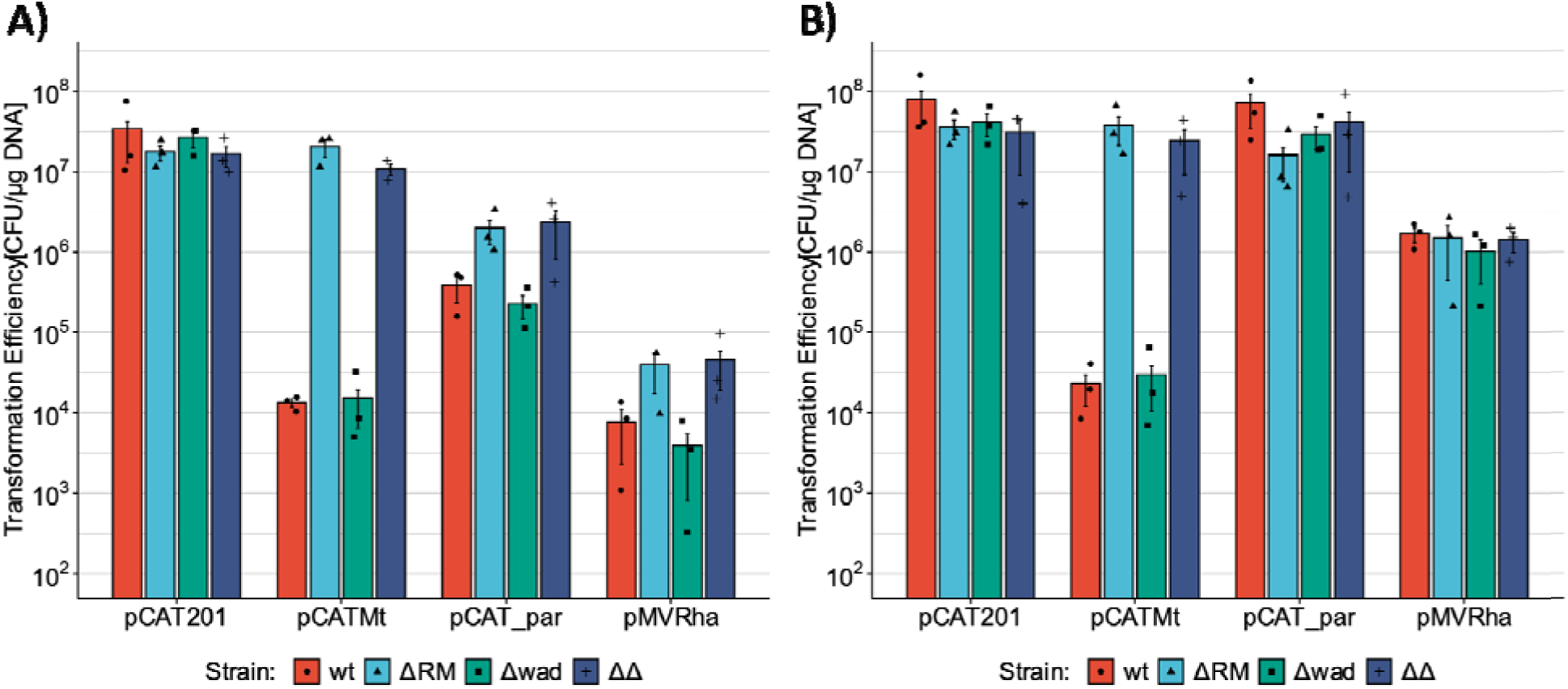
Deletion of the type I and IV RM systems partially restored electroporation efficiency. Transformation efficiency of plasmids pCAT201, pCATMt, pCAT_par and pMVRha in *C. necator* H16, ΔRM, Δwad, ΔΔ; A) Plasmids were extracted from *E. coli* DH5a; B) Plasmids were extracted from *E. coli dam^-^*/*dcm^-^*. Three different batches of competent cells were transformed (mean and standard deviation reported).

Thus, another genetic element present in *C. necator* H16 must be additionally responsible for the selective decrease in transformation efficiency of incorrectly methylated plasmids. The culprit was not the Wadjet system, as deletion did not cause any change in electroporation efficiency compared to the *wt* strain (Figure 5A), and the double knock-out strain *C. necator* ΔΔ performed similarly to the single knock-out *C. necator* ΔRM strain. However, it was reported that this defense system could decrease plasmid stability without affecting electroporation efficiency^59^.

An unstable test plasmid was assembled to investigate the role of the predicted Wadjet system in plasmid instability. This plasmid contained the RSF1010 replicon, which is unstable in *C. necator* when the *par* region is not added^12^. In addition, this plasmid contained an operon that expressed the fluorescent protein GFPmut3 using the J23100 promoter and RBS B0034m^17^. This plasmid was introduced in *C. necator* wt and *C. necator* Δwad by electroporation, and its stability was measured. Under the conditions tested, plasmid RSF1010-GFP was lost completely after about 46 generations (Figure 6) in both strains and no difference was observed between the two genotypes. Therefore, the deleted operon did not affect electroporation efficiency or plasmid stability under the chosen conditions.

**Figure 6:**
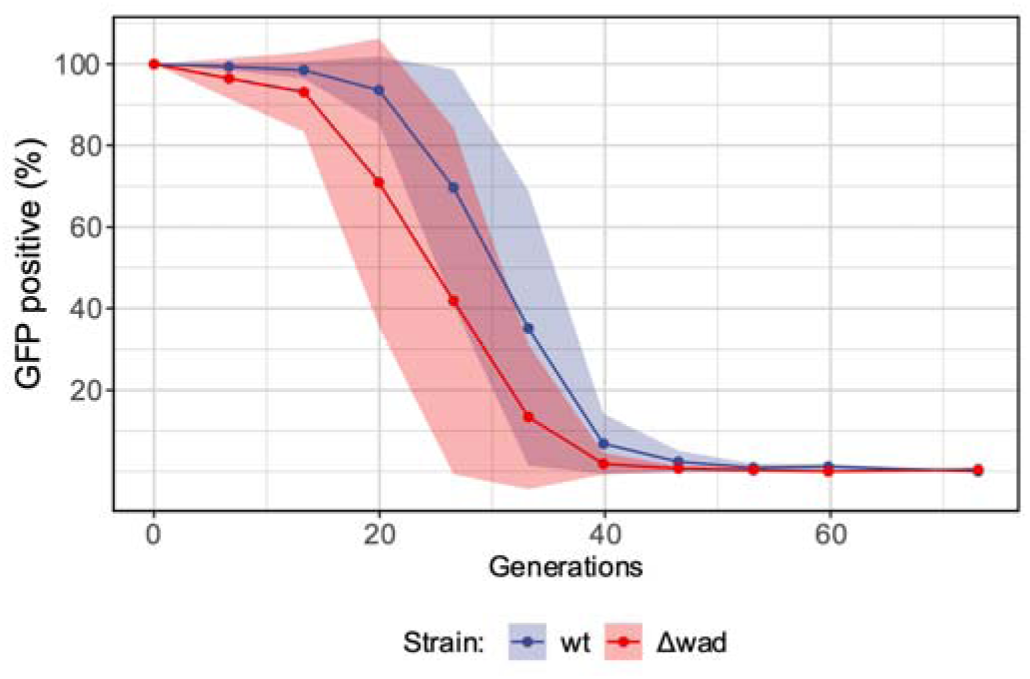
Deletion of the putative Wadjet system did not increase plasmid stability. Plasmid stability of plasmid RSF1010-GFP in *C. necator* wt and *C. necator* Δwad. Two cultures were analyzed for each genotype (mean and standard deviation reported).

## DISCUSSION

Industrial biotechnology has recently shifted its attention from model organisms to a broad range of alternative bacteria, to exploit their unique metabolic capabilities. However, the domestication process is cumbersome and strain-specific. In this study, we characterized the defense systems of the lithoautotrophic bacterium *C. necator* H16 and propose a roadmap to increase electroporation efficiency in other non-model bacteria. *C. necator* has recently been in the spotlight due to its autotrophic metabolism and ability to grow on formate, but its low transformation efficiency hinders extensive genetic modifications. In particular, several research groups have been working towards an increase in electroporation efficiency with promising results. Electroporation protocols have been optimized^12,14^, small plasmids have been designed^12,15^, and putative restriction enzymes have been deleted^16^. These latter deletions increased electroporation efficiency, but the maximum efficiency achieved was still low (about 10^4^ CFU/µg DNA), and thus conjugation is often the method of choice for introducing exogenous DNA.

Therefore, we used *C. necator* as a test case and first compared different electroporation protocols using small and medium size plasmids as standardized test plasmids. We obtained the highest electroporation efficiency ever reported in *C. necator* H16 (10^8^ CFU/µg DNA), but confirmed that restriction avoidance was necessary to introduce larger plasmids. We then proposed a species-independent technique to rapidly clone and express multiple enzymes in novel strains by restriction avoidance. Indeed, using Golden Gate assembly with a natively methylated backbone (here pMVRha) allows rapid and highly efficient cloning and electroporation of multiple inserts in non-model bacteria (70-fold increase in electroporation efficiency in *C. necator*). This technique can be used to express multiple enzymes in newly discovered species to identify the most promising ones. For example, a library of homologs could be screened to identify the best-performing enzyme prior to genomic integration. Large libraries of enzyme variants generated by error-prone PCR could also be rapidly introduced into organisms that are recalcitrant to electroporation. In addition, this technique can be used for other purposes where high transformation efficiency is required. Some Cas9 plasmids, such as the CRISPRi system recently developed for *C. necator* H16, are designed for gRNA cloning using BsaI^63^, and a large number of gRNAs could be cloned and tested simultaneously after purification of the natively methylated backbone. However, there are some limitations. For instance, if the host organism is highly recombinogenic, pMVRha could be mutated before plasmid extraction. In addition, Golden Gate uses BsaI as the restriction enzyme, and if the host organism happens to be able to methylate its recognition pattern, the assembly would not be possible. In this second case, another type IIS restriction enzyme could be used instead.

After this initial screening step, whole genome sequencing using a methylation-sensitive method such as Single Molecule, Real-Time (SMRT) sequencing or Nanopore Sequencing is recommended, and bioinformatic analysis may reveal putative restriction systems, their target patterns and other defense systems^27,28,64^. In *C. necator*, we identified two complete restriction-modification systems with their specificities and a putative Wadjet system that could influence transformation efficiency and plasmid stability in other organisms^29,57–59^. Accordingly, ad-hoc plasmids were created to investigate the role of each predicted system. The plasmids pCATMt, pCAT_par and pMVRha were used to confirm a strong activity of the type I RM system (over 5000-fold decrease of transformation efficiency when its restriction pattern was present), as well as the activity of a type IV restriction system against plasmids larger than pCAT201, with the effect depending on plasmid size (1927-fold decrease for pCAT_par, 5644-fold decrease for pMVRha). We confirmed our bioinformatic predictions and developed a restriction avoidance strategy based on plasmid design (removal of type I RM recognition patterns) and *in vivo* demethylation(using the shuttle strain *E. coli dam-/dcm-*). This greatly increased electroporation efficiency and allowed us to introduce suicide plasmids by electroporation.

Moreover, we successfully deleted both type I and type IV restriction enzymes, and characterized the new strains. These deletions were already described in a previous study^16^. However, our in-depth approach led us to characterize the role of each enzyme, and we clarified a discrepancy in the results observed in a subsequent study (16-fold increase instead of 1697-fold after double deletion) due to the choice of the test plasmid^12^. Surprisingly, we also discovered that a yet undiscovered defense system is still able to target mis-methylated DNA. Furthermore, a putative Wadjet system was identified, but its deletion did not affect either transformation efficiency or plasmid stability under the tested conditions. Indeed, bioinformatic predictions must be treated with caution as they can direct resources to promising targets, but experimental confirmation is always necessary. Many other promising defense systems were identified in *C. necator* H16. A small number of targets were selected for further analysis based on the available literature information. Unfortunately, the role of defense systems in plasmid transformation is yet minimally explored, and thus we speculate that a broader attention to their action against plasmids could benefit the field of industrial biotechnology while shedding light on their sensing and action mechanisms.

## MATERIALS AND METHODS

### Chemicals, bacterial strains, and culture conditions

All chemicals used were purchased from Sigma–Aldrich Ltd, VWR International LLC, or Carl Roth GmbH in the highest purity. *C. necator* H16 (DSM 428) was purchased from DSMZ (Braunschweig, Germany). *E. coli* DH5α was purchased from ThermoFisher Scientific and used as a host for cloning and plasmid propagation. *E. coli dam^-^/dcm^-^* was purchased from New England BioLabs (NEB) and used for de-methylation of plasmid DNA. All microbial strains used are listed in Table 1. *C. necator* and *E. coli* were generally grown in *lysogeny* broth (LB, 10 g/L tryptone, 10 g/L NaCl, 5 g/L yeast extract) or tryptic soy broth (TSB) at 30 °C and 37 °C, respectively. Agar agar was added to a final concentration of 2% to obtain LB agar and TSB agar. When necessary, media was supplemented with antibiotics. Media used for *C. necator* cultivation was supplemented with 20 mg L^-1^ gentamicin, 200 or 400 mg L^-1^ kanamycin, or 15 mg L^-1^ tetracycline. LB supplemented with 400 mg L^-1^ kanamycin was used for selection after electroporation. In all other situations, LB was supplemented with 200 mg L^-1^ kanamycin. Media for *E. coli* DH5α and *E. coli dam^-^/dcm^-^* was supplemented with 50 mg L^-1^ kanamycin or 15 mg L^-1^ tetracycline.

### Cloning and *E. coli* transformation

Plasmid DNA was purified using QIAprep Spin Miniprep Kit (Qiagen) or Monarch® Plasmid Miniprep Kit (NEB). When purifying plasmid DNA from *C. necator*, using less cell culture volume than recommended is advised to avoid genomic DNA contamination (*e.g.* 1.5 - 2 mL instead of 5 mL). DNA purification from PCR reaction mixtures was performed using QIAquick PCR Purification Kit (Qiagen Ltd UK). Microbial genomic DNA was extracted using GenElute Bacterial Genomic DNA Kit (Sigma). DNA was amplified by PCR in 25 μL reactions using Q5® High-Fidelity DNA Polymerase (NEB). All PCR reactions were set up according to the manufacturer’s instructions. BsaI-HF®v2, BbsI-HF, and T4 DNA ligase were purchased from New England BioLabs (NEB). DpnI and T4 Polynucleotide Kinase were purchased from ThermoFisher Scientific.

For *E. coli* transformations, 50 μL of chemically competent cells^65^ were mixed with plasmid DNA, incubated in ice for 30 min, followed by a heat shock at 42 °C for 45 s and a subsequent incubation in ice for 2 min. Cells were recovered in 950 μL of Super Optimal broth with Catabolite repression (SOC) medium at 37 °C for 1 h, plated on LB agar with the appropriate antibiotic and incubated overnight at 37 °C.

### *C. necator* electroporation

Electroporation of *C. necator* was performed according to protocols 1, 2 and 3 derived from Ehsaan *et al.* (2021)^12^, Azubuike *et al.* (2021)^15^ and Tee *et al*. (2017)^14^, respectively. All protocols are described in detail in the Supplementary Information. Protocol 3 was generally used for the preparation of electrocompetent *C. necator* cells and electroporation, unless otherwise stated, with slight modifications to the protocol derived from Tee *et al*.^14^. Briefly, *C. necator* H16 was first streaked onto a TSB agar plate and grown for 40 h. A single colony was then cultivated in SOB supplemented with 20 mg L^-1^ gentamicin for 16 h at 30 °C. Fresh SOB supplemented with gentamicin was inoculated with the preculture at an initial OD_600_ of 0.1 and cultivated at 30 °C. When the cells reached an OD_600_ of 0.4–0.6, they were transferred onto ice and chilled for 5–10 min. The cells were then transferred to 50 mL falcon tubes and centrifuged at 6000 *g* at 4 °C for 2 min. The supernatant was removed, and cells were resuspended in 25 mL of 50 mM CaCl_2_ by briefly using a vortex. They were then incubated for 15 min on ice. The cells were then centrifuged at 6500 *g* at 4 °C for 2 min, and the supernatant was removed. Cells were washed twice using 25 and 15 mL of ice-cold 0.2 M sucrose, respectively. At the end of each wash, cells were centrifuged at 6500 *g* at 4 °C for 2–3 min, and the supernatant was decanted. The cell pellet was finally resuspended in 1/100 of the initial volume (*e.g.*, 100 mL initial cell culture to 1 mL final resuspension volume). 50 μL aliquots of competent cells were transferred into 1.5 mL centrifuge tubes, snap-frozen in liquid nitrogen and stored at −80 °C until further use. For electroporation, each aliquot was thawed on ice for 20 min, transferred into a chilled 1-mm electroporation cuvette, mixed with 50-200 ng of plasmid DNA, incubated for 2-5 min and electroporated (25 μF, 200 Ω, 1.15 kV). 950 μL of SOB supplemented with fructose (20 mM) were immediately added and the cells were transferred to a 2 mL centrifuge tube for outgrowth at 30 °C for 2 h. After the outgrowth, cells were diluted and plated on selective media.

### Plasmid construction

Oligonucleotide primers were synthesized by Eurofins Genomics (Ebersberg, Germany; Supplementary Information, Table S1). Plasmids were sequenced by Sanger sequencing (Macrogen Europe). pCAT201^15^ was purchased from Addgene (#134878) and used as a backbone for further plasmid construction. Plasmid pLO3^42^ was kindly provided by Dr. Oliver Lenz (TU Berlin, Germany). Other genetic parts were PCR-amplified or synthesized from Twist Bioscience. After PCR amplification, template DNA was removed by adding 0.5 μL of DpnI to the PCR mix and incubating the mixture at 37 °C for 1 h. Detailed plasmid construction is described in the Supplementary Information. Plasmids pCAT_par, pMVRha, RSF1010-GFP and pMVRha-GFP were constructed using Golden Gate Assembly^49^. Briefly, for pCAT_par and pMVRha, 75 ng of PCR-amplified backbone were mixed with fragments in a 1:2 molar ratio. For pMVRha-GFP assembly, 75 ng of acceptor plasmid pMVRha were mixed with 75 ng of donor vector c11^52^. For RSF1010-GFP, 75 ng of PCR-amplified backbone was mixed with 75 ng of each donor vector (p13, r03, c11, and te06). Golden Gate reactions were carried out in a total volume of 10 μL by mixing the DNA, MilliQ water, T4 DNA ligase buffer, T4 DNA ligase (200 U), and BsaI-HFv2 or BbsI-HF (6 U). The mixture was then incubated in a thermocycler using the following program: (37°C, 5 min → 16°C, 5 min) x 15 → 60°C, 5 min. Plasmids pLOWad and pLORM were constructed using Gibson assembly^66^. Sequences of pCATMt, pCAT_par and pMVRha are reported in the Supplementary Information.

### Fluorescence measurement

To determine the dose-response of GFP expression from plasmid pMVRha-GFP after rhamnose induction, OD_600_ and GFPmut3 fluorescence were measured over time using a Biolector (M2P Labs, Baesweiler, Germany) in 96 well plates (Greiner Bio-One) with a filling volume of 200 μL. Precultures were cultivated overnight at 30 °C at 200 rpm in glass tubes containing 2 mL of LB supplemented with 200 mg L^-1^ kanamycin. Cultures were then inoculated to an OD_600_ of 0.2 in LB containing varying amounts of L-rhamnose. The Biolector was set to 30 °C, 900 rpm and humidity control of 85 %. Two internal filter modules of the device were used for online measurement. Fluorescence of GFPmut3 was measured at an excitation wavelength of 488 nm and an emission wavelength of 520 nm with a gain 60. Biomass was determined at 620 nm with a gain 40 as scattered light. Scattered light was correlated to OD_600_ with a dilution series of a stationary phase culture. After collecting the data, each value was blanked using a well with empty LB media. To determine specific fluorescence, fluorescence intensity was divided by scattered light. To test whether all the colonies were assembled correctly after Golden Gate assembly, several colonies from each electroporation plate were inoculated in a 96-deep well plate (Enzyscreen) filled with 500 µL LB and kanamycin (200 mg L^-1^) supplemented with 5 mM rhamnose. The plate was then incubated overnight at 30 °C, 300 rpm. 100 µL were then transferred to a 96-well plate (Greiner), and blue light was used to qualitatively observe GFPmut3 fluorescence.

### Gene deletion

Knock-out plasmids pLOWad and pLORM were first transferred in *E. coli dam-/dcm-* to remove DNA methylation. The purified plasmids were then used for *C. necator* H16 electroporation. Transconjugants were selected on LB supplemented with 15 mg L^-1^ tetracycline. 1-4 positive colonies were picked and grown overnight in 5 mL of low salt LB (LSLB) without any supplementation (10 g L^-1^ tryptone, 5 g L^-1^ NaCl, 5 g L^-1^ yeast extract). Dilutions were made, and 100 μL of each dilution were plated on LSLB-agar supplemented with 150 g L^-1^ sucrose and incubated at 30 °C for 48 h to select for *sacB* negative colonies (i.e., double recombination). Colony PCR was performed on sucrose-resistant colonies using a flanking pair of oligonucleotide primers to screen for the double crossover deletion mutants. Promising clones were then grown overnight for genomic DNA extraction. Flanking primers were used to amplify the targeted region, and the amplicon was sequenced to confirm deletion (Supplementary Information, Figures S2 and S3).

### Plasmid stability

Single colonies were inoculated in 2 mL of LB supplemented with 200 mg L^-1^ kanamycin and grown overnight at 30 °C, 200 rpm. The cultures were then diluted 1:100 in fresh LB without antibiotics and grown for 24 h. The dilution was then repeated each day. Every day, OD_600_ of each culture was measured, and the cultures were serially diluted and plated in LB agar without supplementation. After 3 days, colonies were counted. GFP-positive colonies were identified using blue light. The ratio of GFP-positive colonies and total colonies was used to determine plasmid presence. Generation number *n* was calculated using the following formula: 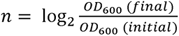

## AUTHOR INFORMATION

### Corresponding Author

Sandy Schmidt − Department of Chemical and Pharmaceutical Biology, University of Groningen, Groningen Research Institute of Pharmacy, Antonius Deusinglaan 1, 9713AV Groningen, The Netherlands; orcid.org/0000-0002-8443-8805;

### Authors

**Matteo Vajente** − Department of Chemical and Pharmaceutical Biology, University of Groningen, Groningen Research Institute of Pharmacy, Antonius Deusinglaan 1, 9713AV Groningen, The Netherlands; orcid.org/0000-0002-8589-9072;

**Riccardo Clerici** − Institute of Applied Microbiology (iAMB), Aachen Biology and Biotechnology (ABBt), RWTH Aachen University, Worringerweg 1, 52074 Aachen, Germany; orcid.org/ 0009-0009-9721-1544;

**Hendrik Ballerstedt** − Institute of Applied Microbiology (iAMB), Aachen Biology and Biotechnology (ABBt), RWTH Aachen University, Worringerweg 1, 52074 Aachen, Germany; https://orcid.org/0000-0001-5729-1724;

**Lars M. Blank** − Institute of Applied Microbiology (iAMB), Aachen Biology and Biotechnology (ABBt), RWTH Aachen University, Worringerweg 1, 52074 Aachen, Germany; https://orcid.org/0000-0003-0961-4976;

## ACKNOWLEDGEMENTS

This project has received funding from the European Union’s Horizon 2020 research and innovation programme under the Marie Skłodowska Curie grant agreement No 955740. S.S. also acknowledges financial support from the Dutch Research Council (grant no. OCENW.XS22.1.044). The laboratory of LMB is partially funded by the Deutsche Forschungsgemeinschaft (DFG, German Research Foundation) under Germany’s Excellence Strategy—Cluster of Excellence 2186 “The Fuel Science Center”—ID: 390919832.

