## Supplementary for "Using *Cupriavidus necator* H16 to provide a roadmap for increasing electroporation efficiency in non-model bacteria"

### SUPPLEMENTARY INFORMATION

#### 1. Experimental section

##### 1. Plasmid construction

pCAT_par was constructed using Golden Gate assembly (full sequence provided below). Oligonucleotide primers pCATpar_back_fw/rev were used to amplify the backbone from plasmid pCAT201[1] (Addgene, #134878). Oligonucleotides pCATpar_par_fw/rev were used to amplify the partitioning region from plasmid pKRC[2,3], kindly provided by Dr. Petra Heidinger (TU Graz/acib, Austria). Golden Gate assembly was executed as described using BsaI-HFv2. Correct assembly was confirmed by Sanger sequencing.

pMVRha was constructed using Golden Gate assembly (full sequence provided below). Oligonucleotide primers pMVRha_back_fw/rev were used to amplify the backbone from plasmid pMVCu (unpublished). Oligonucleotides pMVRha_Rha_fw/rev were used to amplify the rhamnose inducible system from pKRrha (kindly provided by Prof. Dirk Holtmann, KIT, Germany)[4]. Golden Gate assembly was executed as described using BbsI-HF. pMVRha contains a backbone formed by pBBR1 origin of replication, RP4 mobilization sequence (*oriT*), kanamycin resistance marker and rhamnose inducible system. Two BsaI cutting sites flank an operon constituted by a strong promoter ($P_{j5[c2]}$,[5]) which drives the expression of cloning marker *spisPink*[6]. Correct assembly was confirmed by Sanger sequencing.

pCATMt was constructed using Golden Gate assembly (full sequence provided below). Oligonucleotide primers pCATMt_back_fw/rev were used to amplify the backbone from plasmid pCAT201. Oligonucleotides pCATMt_ins_fw/rev were annealed to obtain dsDNA. Shortly, 9 µL of each primer (100 µM) were mixed with annealing buffer in a final volume of 30 µL (final concentration: 10 mM Tris; 50 mM NaCl; 1 mM EDTA). The mix was incubated in a thermocycler using the following program: 95 °C, 2 min → (95 °C, 40 s) x 70 cycles (temperature decreases by 1 °C each cycle). 0.3 µL of this mixture were then phosphorylated using T4 Polynucleotide Kinase (ThermoFisher) according to the manufacturer's instructions. Subsequently, Golden Gate assembly was executed using 75 ng of backbone and a 1:10 molar ratio of dsDNA phosphorylated insert. Correct assembly was confirmed by Sanger sequencing.

pLO3Mt was constructed by inserting a silent mutation in pLO3 to remove a *C. necator* restriction site (GAYNNNNNCTTGY) by site-directed mutagenesis[7]. Shortly, oligonucleotide primers pLO3Mt_ fw/rev were used to amplify the plasmid pLO3 (kindly provided by Dr. Oliver Lenz (TU Berlin, Germany))[8] and to introduce the silent mutation. Template DNA was removed by DpnI digestion. The two DNA molecules were mixed and *E. coli* DH5α was then transformed with 2 µL of the mixture. The successful mutation was confirmed by Sanger sequencing.

pLOWad was constructed using Gibson assembly. Oligonucleotide primers pLOWad_pLO_fw/rev were used to amplify the backbone from plasmid pLO3Mt. Oligonucleotides pLOWad_UP_fw/rev and pLOWad_DOWN_fw/rev were used to amplify two 1 kb regions from *C. necator* genomic DNA. The successful assembly was confirmed by Sanger sequencing.

pLORM was constructed using Gibson assembly. Oligonucleotide primers pLORM_pLO_fw/rev were used to amplify the backbone from plasmid pLO3Mt. Oligonucleotides pLORM_UP_fw/rev and pLORM_DOWN_fw/rev were used to amplify two 1 kb regions from *C. necator* genomic DNA. The successful assembly was confirmed by Sanger sequencing.

RSF1010-GFP was constructed using Golden Gate assembly. Shortly, primers RSF1010_back_fw/rev were used to amplify the backbone from plasmid pSEVA251[9]. Template was removed by DpnI digestion. Plasmids p13, r03, c11, and te06 were rehydrated and extracted from the Golden Standard library[10]. Golden Gate assembly was executed as described using BsaI-HF. The correct assembly was confirmed by Sanger sequencing.

#### 2. Preparation of electrocompetent *C. necator* cells and electroporation protocols

Electroporation of *C. necator* was performed according to protocols 1, 2 and 3 derived from Ehsaan *et al.*[11], Azubuike *et al.*[1] and Tee *et al.*[12], respectively.

*Preparation and transformation of electrocompetent C. necator cells according to protocol 1:*

First, *C. necator* H16 was streaked on a TSB agar plate and grown at 30 °C for 40 h. A heavy loop of freshly grown *C. necator* cells was inoculated in 10 mL of SOB supplemented with 20 mg $L^{-1}$ gentamicin and resuspended thoroughly. $10^{-1}$, $10^{-2}$ and $10^{-3}$ serial dilutions were prepared in SOB supplemented with 20 mg $L^{-1}$ gentamicin and grown overnight at 30 °C, 250 rpm. Fresh SOB supplemented with 20 mg $L^{-1}$ gentamicin was then inoculated with a culture in mid- or late-logarithmic phase at an initial optical density at 600 nm ($OD_{600}$) of 0.055-0.075 and incubated at 30 °C until the culture reached an $OD_{600}$ of 0.25-0.3. Cells were transferred to 50 mL centrifuge tubes, 25 mL in each tube. Biomass was pelleted by centrifugation (5369 *g*, 10 min, 4 °C) and the supernatant was discarded. Cells were resuspended in 10 mL of pre-chilled buffer A (1 mM HEPES, adjusted to pH 7.0 with NaOH, filter-sterilized) and centrifuged as before. The supernatant was discarded and cells were washed with 5 mL of buffer A as described before. Cells were then re-suspended in buffer A supplemented with 10% glycerol (*w/v*) to a final $OD_{600}$ of 5. 100 µL aliquots of competent cells were transferred into 1.5 mL centrifuge tubes, snap-frozen in liquid nitrogen and stored at -80 °C until needed. For electroporation, 100 µL of electro-competent cells were thawed, transferred to a pre-chilled 1-mm electroporation cuvette, mixed with 50-100 ng of plasmid DNA, incubated for 2-5 min and electroporated (25 µF, 200 Ω, 1.25 kV). 950 µL of SOB supplemented with fructose (20 mM) were immediately added and the cells were then transferred to a 2 mL centrifuge tube for outgrowth at 30 °C for 2 h. After the outgrowth, cells were diluted and plated on selective media.

*Preparation and transformation of electrocompetent C. necator cells according to protocol 2:*

First, *C. necator* H16 was streaked on a TSB agar plate and grown at 30 °C for 40 h. A single colony was cultivated in SOB supplemented with 20 mg $L^{-1}$ gentamicin for 16 h at 28 °C. For electrocompetent cell preparation, fresh SOB supplemented with gentamicin was inoculated with the preculture at a 1:1000 dilution and cultivated at 30 °C. When the cells reached an $OD_{600}$ of 0.4–0.8, they were centrifuged at 986 *g*, 4 °C for 10 min. The supernatant was removed, and cells were washed twice with 50 mL of ice-cold 10% (w/v) glycerol. After the last centrifugation, the cells were resuspended in 1/100 of the initial volume (e.g., 100 mL initial cell culture to 1 mL final resuspension volume). 50 µL aliquots of competent cells were transferred into 1.5 mL centrifuge tubes, snap-frozen in liquid nitrogen and stored at -80 °C until needed. For electroporation, each aliquot was thawed on ice, transferred into a chilled 1 mm electroporation cuvette, mixed with 50-200 ng of plasmid DNA, incubated for 2-5 min and electroporated (25 µF, 200 Ω, 1.25 kV). 950 µL of SOB supplemented with fructose (20 mM) were immediately added and the cells were then transferred to a 2 mL centrifuge tube for outgrowth at 30°C for 2 h. After the outgrowth, cells were diluted and plated on selective media.

*Preparation and transformation of electrocompetent C. necator cells according to protocol 3:*

Protocol 3 was generally used for the preparation of electrocompetent *C. necator* cells and electroporation, unless otherwise stated, with slight modifications to the protocol derived from Tee *et al.*[12]. Briefly, *C. necator* H16 was first streaked onto a TSB agar plate and grown for 40 h. A single colony was then cultivated in SOB supplemented with 20 mg $L^{-1}$ gentamicin for 16 h at 30 °C. Fresh SOB supplemented with gentamicin was inoculated with the preculture at an initial $OD_{600}$ of 0.1 and cultivated at 30 °C. When the cells reached an $OD_{600}$ of 0.4–0.6, they were transferred onto ice and chilled for 5–10 min. The cells were then transferred to 50 mL falcon tubes and centrifuged at 6000 *g* at 4 °C for 2 min. The supernatant was removed, and cells were resuspended in 25 mL of 50 mM $CaCl_2$ by briefly using a vortex. They were then incubated for 15 min on ice. The cells were then centrifuged at 6500 *g* at 4 °C for 2 min and the supernatant was removed. Cells were washed twice using 25 and 15 mL of ice-cold 0.2 M sucrose, respectively. At the end of each wash, cells were centrifuged at 6500 *g* at 4 °C for 2–3 min and the supernatant was decanted. The cell pellet was finally resuspended in 1/100 of the initial volume (*e.g.*, 100 mL initial cell culture to 1 mL final resuspension volume). 50 µL aliquots of competent cells were transferred into 1.5 mL centrifuge tubes, snap-frozen in liquid nitrogen and stored at -80 °C until further use. For electroporation, each aliquot was thawed on ice for 20 min, transferred into a chilled 1-mm electroporation cuvette, mixed with 50-200 ng of plasmid DNA, incubated for 2-5 min and electroporated (25 µF, 200 Ω, 1.15 kV). 950 µL of SOB supplemented with fructose (20 mM) were immediately added and the cells were transferred to a 2 mL centrifuge tube for outgrowth at 30 °C for 2 h. After the outgrowth, cells were diluted and plated on selective media.

#### 2. Supplementary Tables

***Table S1:*** *List of putative defense systems identified by PADLOC*[13]*, DefenseFinder*[14] *and REBASE* [15]*. In "Identified by": DF: DefenseFinder, RE: REBASE, PD: PADLOC. The putative methylated residues are bold. Genomic sequence from NCBI (RefSeq assembly accession: GCF_004798725.1).*

| **Defense system** | **Identified by:** | **Gene(s)** | **Reference** |
|---|---|---|---|
| Restriction-Modification system type I | PD, RE, DF | E6A55_RS00020 (Methyltransferase subunit);<br>E6A55_RS00025 (Specificity subunit);<br>E6A55_RS00030 (Restriction subunit) | Loenen *et al.*, 2014[16]<br>In REBASE, the predicted recognition pattern of this system is G**A**YNNNNNC**T**TGY. |
| Restriction system type IV | RE | E6A55_RS00040;<br>E6A55_RS00045 | Loenen and Raleigh, 2014[17] |
| Orphan methyltransferase | RE | E6A55_RS33850 | In REBASE, the predicted recognition pattern of this enzyme is G**T**WW**A**C. |
| Wadjet type III | PD, DF | E6A55_RS00090 (JetD3);<br>E6A55_RS00095 (JetC3);<br>E6A55_RS00100 (JetB3);<br>E6A55_RS00105 (JetA3) | Panas *et al.*, 2014; Doron *et al.*, 2018; Deep *et al.*, 2022; Liu *et al.*, 2022[18–21] |
| DRT class III | PD | E6A55_RS04470 (Drt1a) | Gao *et al.*, 2020[22] |
| AbiE | PD | E6A55_RS09410 (pseudo);<br>E6A55_RS09425 (pseudo) | Dy *et al.*, 2014[23] |
| retron_I-B | PD | E6A55_RS12740 (ATPase-Toprim I-B);<br>E6A55_RS12745 (RT I-B);<br>msr-msd | Gao *et al.*, 2020; Millman *et al.*, 2020[22,24] |
| Zorya type III | PD | E6A55_RS21755 (ZorF3);<br>E6A55_RS21760 (ZorB3);<br>E6A55_RS21765 (ZorA3);<br>E6A55_RS21770 (ZorG3) | Doron *et al.*, 2018[19] |
| AVAST type I | PD, DF | E6A55_RS32990 (Avs1c);<br>E6A55_RS32995 (Avs1b);<br>E6A55_RS33000 (Avs1a) | Gao *et al.*, 2020[22] |
| Gabija | PD, DF | E6A55_RS33010 (GajB);<br>E6A55_RS33015 (GajA) | Doron *et al.*, 2018[19] |
| MazEF | DF | E6A55_RS34165;<br>E6A55_RS34160 | Nikolic *et al.*, 2023[25] |
| dGTPase | PD | E6A55_RS17515 | Tal *et al.*, 2022[26] |
| ietAS | PD | E6A55_RS32710 (IetA);<br>E6A55_RS32715 (IetS) | Gao *et al.*, 2020[22] |
| PD-T4-6 | PD | E6A55_RS04480;<br>E6A55_RS04815 | Vassallo *et al.*, 2022[27] |

***Table S2**: Oligonucleotides used in this work.*

| **Primer** | **Sequence** |
|---|---|
| pCATpar_back_fw* | AAGGGTCTCATGCCAGCAAGCCCGTAGGG |
| pCATpar_back_rev* | AAGGGTCTCAGCAATGCTCTCCGGGCTTC |
| pCATpar_par_fw* | AAGGGTCTCATTGCGCGAAAAGGTGAGAAAAGCC |
| pCATpar_par_rev* | AAGGGTCTCAGGCAAGGGCATGAAAAAGCCCGT |
| pMVRha_back_fw* | AGAAGACAAATGTGAGACCCGCAGAAAG |
| pMVRha_back_rev* | AGAAGACAAGCAAAAAACCCCTCAAGACC |
| pMVRha_Rha_fw* | AGAAGACAATTGCTTAATCTTTCTGCGAATTGAG |
| pMVRha_Rha_rev* | AGAAGACAAACATTTGTATATCTCCTTCTTAAGAATTG |
| pCATMt_back_fw* | AAGGTCTCAGTTAATTAAGTTCCAGACAAG |
| pCATMt_back_rev* | AAGGTCTCACCTATTGGTTAAAAAATGAGC |
| pCATMt_ins_fw | TAGGGCAAGCAGGCATC |
| pCATMt_ins_rev | TAACGATGCCTGCTTGC |
| pLO3Mt_ fw | AAGAAAACAAGCGTTGTCAAAGACAGCATCCTTGAACAAGGAC |
| pLO3Mt_ rev | TTGACAACGCTTGTTTTCTTGCCTTTGATGTTCAGCAGGAAGC |
| pLOWad_pLO_fw | CGATCCACCCGGCCGATTTCCTGCAGGCATGCAAGCTAATTC |
| pLOWad_pLO_rev | CCGCCATAAGCGGCCGTCGAGAGCTCGAATTAAAAGGATCTAGG |
| pLOWad_UP_fw | GATCCTTTTAATTCGAGCTCTCGACGGCCGCTTATGGCGG |
| pLOWad_UP_rev | GCTTCCTTTTGCGATCCCGCAGCGCGGAAATGGCGGCCTC |
| pLOWad_DOWN_fw | GAGGCCGCCATTTCCGCGCTGCGGGATCGCAAAAGGAAGC |
| pLOWad_DOWN_rev | ATTAGCTTGCATGCCTGCAGGAAATCGGCCGGGTGGATCG |
| pLORM_pLO_fw | CTGCTGACGGGTTGGCAATCTGCAGGCATGCAAGCTAATTC |
| pLORM_pLO_rev | CGGACGAAAAACACTGCTCGAGCTCGAATTAAAAGGATCTAGG |
| pLORM_UP_fw | GATCCTTTTAATTCGAGCTCGAGCAGTGTTTTTCGTCCGG |
| pLORM_UP_rev | ACTCATCCGATCAGACTTGGTCAGGCGCTCCCTGCTTG |
| pLORM_DOWN_fw | AACAAGCAGGGAGCGCCTGACCAAGTCTGATCGGATGAGTC |
| pLORM_DOWN_rev | AATTAGCTTGCATGCCTGCAGATTGCCAACCCGTCAGCAG |
| RSF1010_back_fw* | AGGTCTCACTCCGTCGTGACTGGGAAAACCC |
| RSF1010_back_rev* | AGGTCTCACGCTCCTGTGTGAAATTGTTATCCGC |
| RM _1_fw | GAAGGGTAAGGCACCACTCG |
| RM _1_rev | TCTACCAGCGGCACAGTTCC |
| Wad_1_fw | CGGGATAGCTGGTCCATGAC |
| Wad_1_rev | AGCTTATGACGGCCAAGGAC |

**Golden Gate enzyme recognition site is underlined. Golden Gate enzyme restriction site is highlighted in purple.*

#### 3. Supplementary Figures

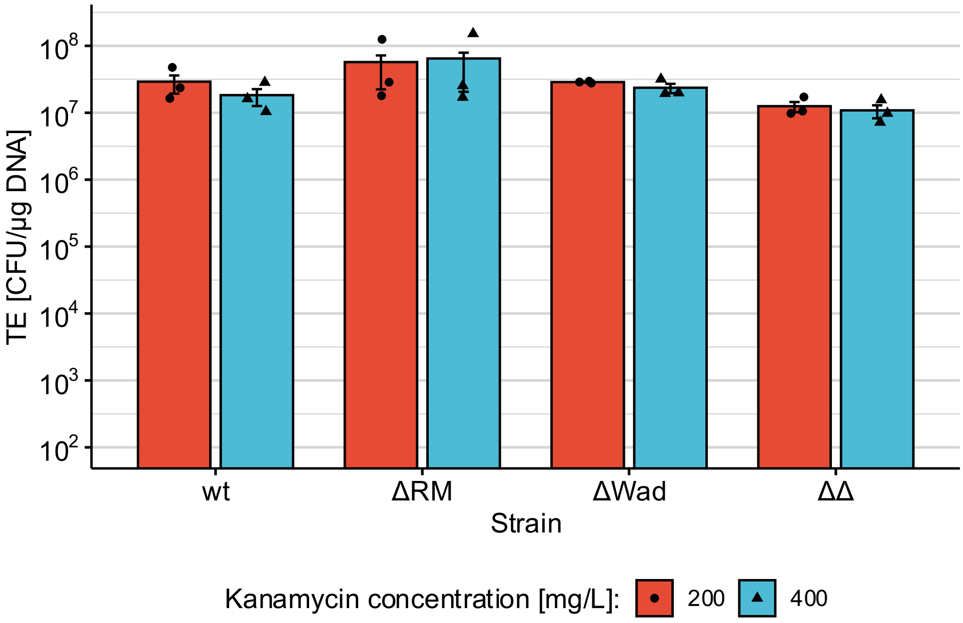


**Figure S1**: Electroporation efficiency of different *C. necator* strains transformed with pCAT201. After outgrowth, cells were plated in LB agar supplemented with 200 and 400 mg $L^{-1}$ of kanamycin and grown for 40 h at 30 °C. For each condition tested, three transformations were performed (mean and standard deviation reported).

A)

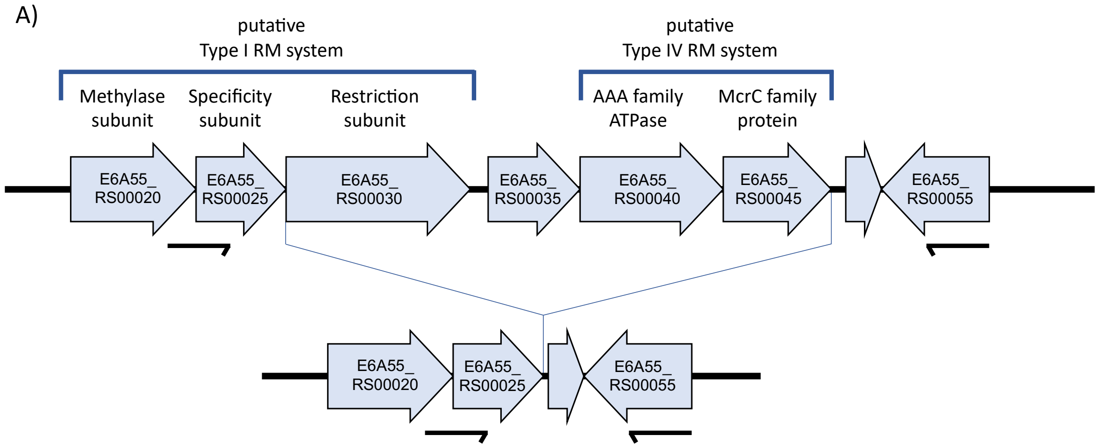


B)

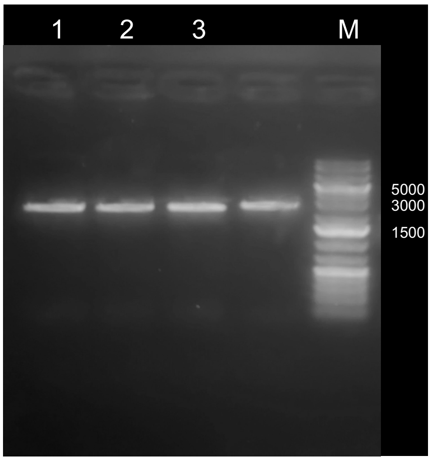


**Figure S2**: Deletion of genes E6A55_RS00030 - E6A55_RS00045 (H16_A0006 - H16_A0009). A) Schematic representation of the wild type and putative mutant genomes in the targeted region, together with the position of the primers used (RM _1_fw/rev). B) Genomic DNA of promising colonies was extracted. Genomic DNA was then amplified using Q5 polymerase. Electropherogram of the PCR products: M = GeneRuler 1 kb plus DNA ladder (ThermoFisher). Lanes 1-3, PCR of putative knock-out strains using primers RM_1_fw/rev (expected amplicon size: 2498 bp). The amplified fragment was then sequenced to confirm successful deletion.

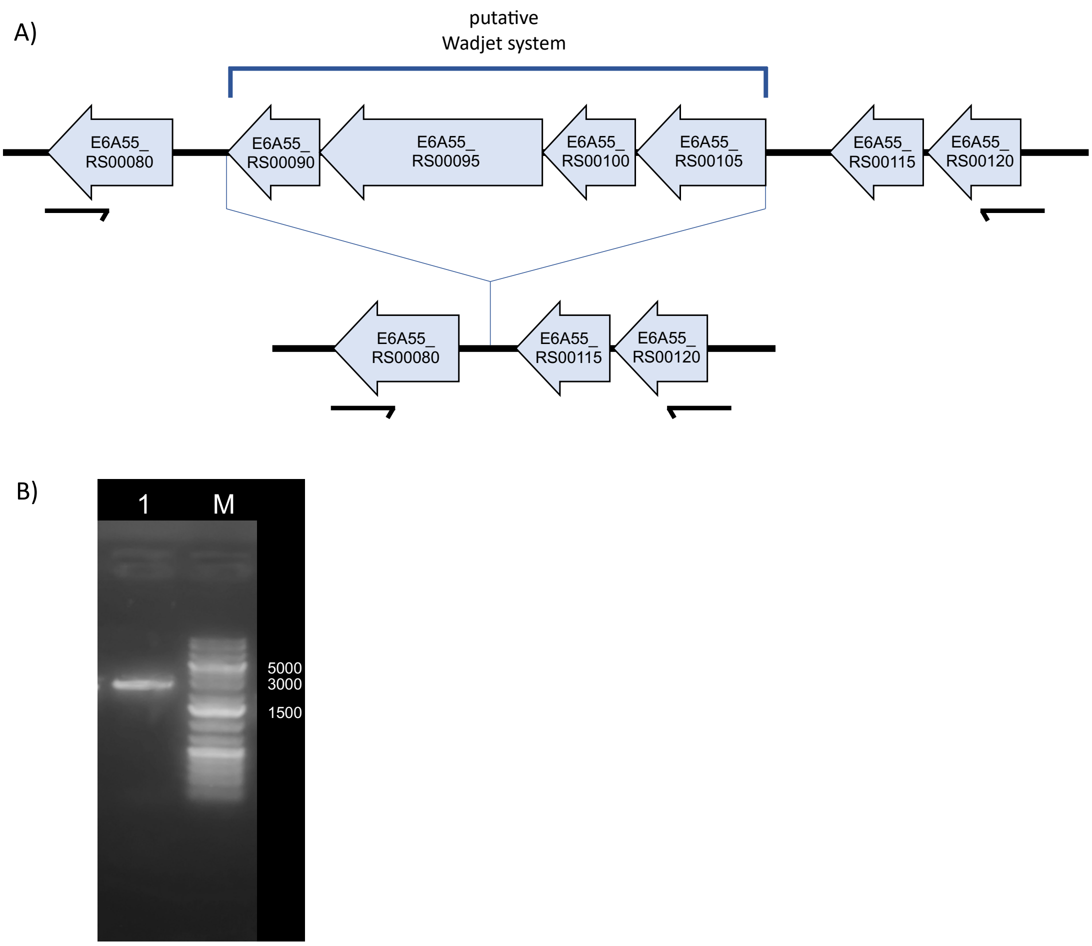


**Figure S3**: Deletion of E6A55_RS00090 - E6A55_RS00105 (H16_A0017 - H16_A0020). A) Schematic representation of the wild type and putative mutant genomes in the targeted region, together with the position of the primers used (Wad_1_fw/rev). B) Genomic DNA of promising colonies was extracted and amplified using Q5 polymerase. Electropherogram of the PCR products: M = GeneRuler 1 kb plus DNA ladder (ThermoFisher). Lane 1, PCR of putative knock-out strain using primers Wad_1_fw/rev (expected amplicon size: 2600 bp). The amplified fragment was then sequenced to confirm successful deletion.

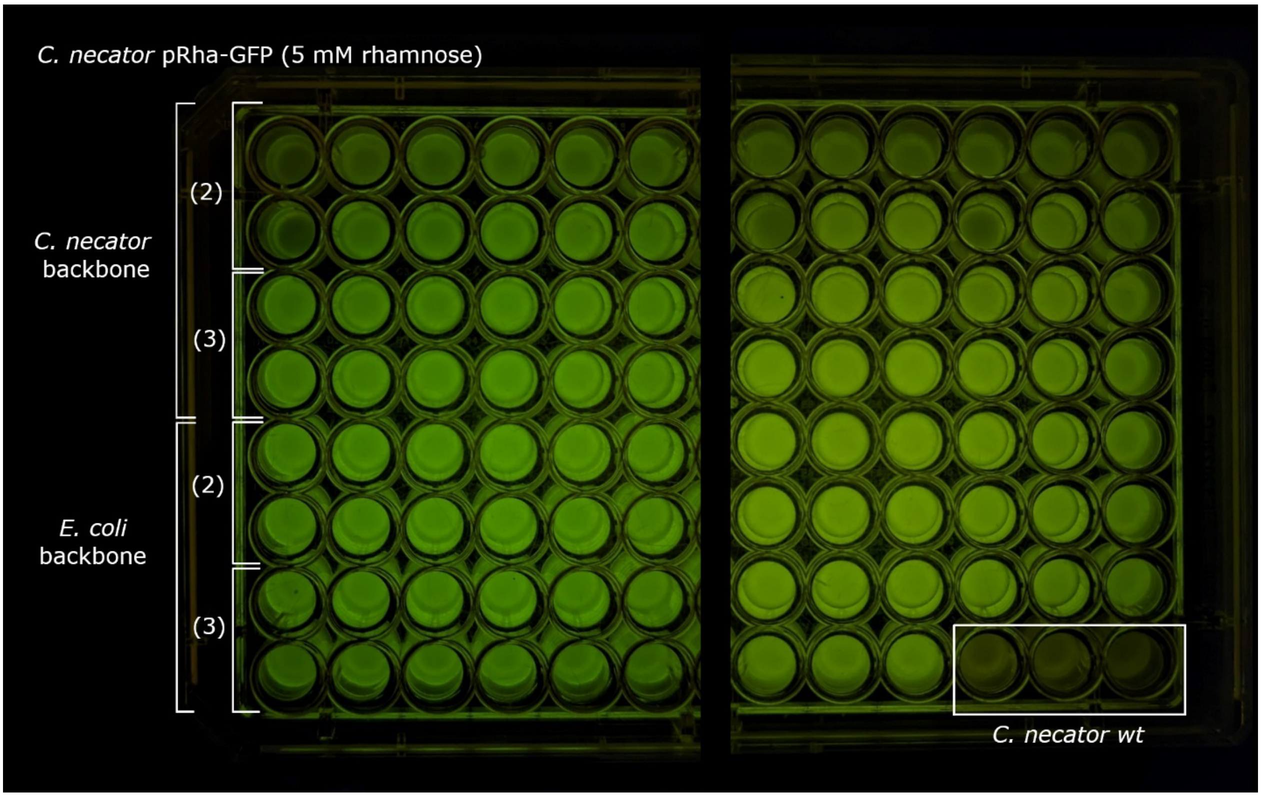


**Figure S4**: GFPmut3 expression of randomly picked colonies after one step Golden Gate assembly and electroporation of pMVRha-GFP. 48 colonies were picked from pMVRha-GFP assembled with backbone extracted from *C. necator*. 45 colonies were picked from pMVRha-GFP assembled with backbone extracted from E. coli. Three wells were inoculated with C. necator wt (bottom right, negative control). Shortly, colonies were picked and inoculated in a 96-deep well plate, each well containing 0,5 mL of LB supplemented with rhamnose (5 mM). The plate was incubated overnight at 30 °C, 300 rpm. 100 µL from each well were then transferred to a clear 96-well plate (Greiner) and imaged.

#### 4. Plasmid sequences

**Sequence of plasmid pCATMt (3200 bp):**

acgctgacttgacgggacggcgcgccgcttcactcattaggcaccccaggctttacactttatgcttccggctcgtatgttgtgtggaattgtgagcggataaca
atttcacacaggaaacagctatgaccatgattacgccaagcgcgcaattaaccctcactaaagggaacaaaagctgggtaccgggccccccctcgaggtcg
acggtatcgataagcttgatatcgaattcctgcagcccgggggatccactagttctagagcggccgccaccgcggtggagctccaattcgccctatagtgagt
cgtattacgcgcgctcactggccgtcgttttacaacgtcgtgactgggaaaaccctggcgttacccaacttaatcgccttgcagcacatccccctttcgccagct
ggcgtaatagcgaagaggcccgcaccgatcgcccttcccaacagttgcgcagcctgaatggcgaatggaaattgtaagcgttaatattttgttaaaattcgcgtt
aaatttttgttaaatcagctcattttttaaccaatagggcaagcaggcatcgttaattaagttccagacaaggtatagggcggcgaggcggctacagccgatagt
ctggaacagcgcacttacgggttgctgcgcaacccaagtgctaccggcgcggcagcgtgacccgtgtcggcggctccaacggctcgccatcgtccagaaa
acacggctcatcgggcatcggcaggcgctgctgcccgcgccgttcccattcctccgtttcggtcaaggctggcaggtctggttccatgcccggaatgccggg
ctggctgggcggctcctcgccggggccggtcggtagttgctgctcgcccggatacagggtcgggatgcggcgcaggtcgccatgccccaacagcgattcg
tcctggtcgtcgtgatcaaccaccacggcggcactgaacaccgacaggcgcaactggtcgcggggctggccccacgccacgcggtcattgaccacgtagg
ccgacacggtgccggggccgttgagcttcacgacggagatccagcgctcggccaccaagtccttgactgcgtattggaccgtccgcaaagaacgtccgatg
agcttggaaagtgtcttctggctgaccaccacggcgttctggtggcccatctgcgccacgaggtgatgcagcagcattgccgccgtgggtttcctcgcaataa
gcccggcccacgcctcatgcgctttgcgttccgtttgcacccagtgaccgggcttgttcttggcttgaatgccgatttctctggactgcgtggccatgcttatctcc
atgcggtaggggtgccgcacggttgcggcaccatgcgcaatcagctgcaacttttcggcagcgcgacaacaattatgcgttgcgtaaaagtggcagtcaatta
cagattttctttaacctacgcaatgagctattgcggggggtgccgcaatgagctgttgcgtaccccccttttttaagttgttgatttttaagtctttcgcatttcgcccta
tatctagttctttggtgcccaaagaagggcacccctgcggggttcccccacgccttcggcgcggctccccctccggcaaaaagtggcccctccggggcttgtt
gatcgactgcgcggccttcggccttgcccaaggtggcgctgccccttggaacccccgcactcgccgccgtgaggctcgggggcaggcgggcgggctt
cgcccttcgactgcccccactcgcataggcttgggtcgttccaggcgcgtcaaggccaagccgctgcgcggtcgctgcgcgagccttgacccgccttccact
tggtgtccaaccggcaagcgaagcgcgcaggccgcaggccggaggcttttccccagagaaaattaaaaaaattgatggggcaaggccgcaggccgcgca
gttggagccggtgggtatgtggtcgaaggctgggtagccggtgggcaatccctgtggtcaagctcgtgggcaggcgcagcctgtccatcagcttgtccagc
agggttgtccacgggccgagcgaagcgagccagccggtggccgctcgcggccatcgtccacatatccacgggctggcaagggagcgcagcgaccgcg
cagggcgaagcccggagagcaagcccgtagggcgccgcagccgccgtaggcggtcacgactttgcgaagggccggccacctggaaagccacgttgtgt
ctcaaaatctctgatgttacattgcacaagataaaaatatatcatcatgaacaataaaactgtctgcttacataaacagtaatacaaggggtgttatgagccatattc
aacgggaaacgtcttgctcgaggccgcgattaaattccaacatggatgctgatttatatgggtataaatgggctcgcgataatgtcgggcaatcaggtgcgaca
atctatcgattgtatgggaagcccgatgcgccagagttgtttctgaaacatggcaaaggtagcgttgccaatgatgttacagatgagatggtcagactaaactgg
ctgacggaatttatgcctcttccgaccatcaagcattttatccgtactcctgatgatgcatggttactcaccactgcgatccccgggaaaacagcattccaggtatt
agaagaatatcctgattcaggtgaaaatattgttgatgcgctggcagtgttcctgcgccggttgcattcgattcctgtttgtaattgtccttttaacagcgatcgcgta
tttcgtctcgctcaggcgcaatcacgaatgaataacggtttggttgatgcgagtgattttgatgacgagcgtaatggctggcctgttgaacaagtctggaaagaa
atgcataagcttttgccattctcaccggattcagtcgtcactcatggtgatttctcacttgataaccttatttttgacgaggggaaattaataggttgtattgatgttgga
cgagtcggaatcgcagaccgataccaggatcttgccatcctatggaactgcctcggtgagttttctccttcattacagaaacggctttttcaaaaatatggtattga
taatcctgatatgaataaattgcagtttcatttgatgctcgatgagtttttctaatcagaattggttaattggttgtaacactggcagagcatt

**Sequence of plasmid pCAT_par (5448 bp):**

acgctgacttgacgggacggcgcgccgcttcactcattaggcaccccaggctttacactttatgcttccggctcgtatgttgtgtggaattgtgagcggataaca
atttcacacaggaaacagctatgaccatgattacgccaagcgcgcaattaaccctcactaaagggaacaaaagctgggtaccgggccccccctcgaggtcg
acggtatcgataagcttgatatcgaattcctgcagcccgggggatccactagttctagagcggccgccaccgcggtggagctccaattcgccctatagtgagt
cgtattacgcgcgctcactggccgtcgttttacaacgtcgtgactgggaaaaccctggcgttacccaacttaatcgccttgcagcacatccccctttcgccagct
ggcgtaatagcgaagaggcccgcaccgatcgcccttcccaacagttgcgcagcctgaatggcgaatggaaattgtaagcgttaatattttgttaaaattcgcgtt
aaatttttgttaaatcagctcattttttaaccaataggccgactgcgatgagttaattaagttccagacaaggtatagggcggcgaggcggctacagccgatagtc
tggaacagcgcacttacgggttgctgcgcaacccaagtgctaccggcgcggcagcgtgacccgtgtcggcggctccaacggctcgccatcgtccagaaaa
cacggctcatcgggcatcggcaggcgctgctgcccgcgccgttcccattcctccgtttcggtcaaggctggcaggtctggttccatgcccggaatgccgggc
tggctgggcggctcctcgccggggccggtcggtagttgctgctcgcccggatacagggtcgggatgcggcgcaggtcgccatgccccaacagcgattcgt
cctggtcgtcgtgatcaaccaccacggcggcactgaacaccgacaggcgcaactggtcgcggggctggccccacgccacgcggtcattgaccacgtagg
ccgacacggtgccggggccgttgagcttcacgacggagatccagcgctcggccaccaagtccttgactgcgtattggaccgtccgcaaagaacgtccgatg
agcttggaaagtgtcttctggctgaccaccacggcgttctggtggcccatctgcgccacgaggtgatgcagcagcattgccgccgtgggtttcctcgcaataa
gcccggcccacgcctcatgcgctttgcgttccgtttgcacccagtgaccgggcttgttcttggcttgaatgccgatttctctggactgcgtggccatgcttatctcc

atgcggtaggggtgccgcacggttgcggcaccatgcgcaatcagctgcaacttttcggcagcgcgacaacaattatgcgttgcgtaaaagtggcagtcaatta
cagattttctttaacctacgcaatgagctattgcggggggtgccgcaatgagctgttgcgtaccccccttttttaagttgttgatttttaagtctttcgcatttcgcccta
tatctagttctttggtgcccaaagaagggcacccctgcggggttcccccacgccttcggcgcggctccccctccggcaaaaagtggcccctccggggcttgtt
gatcgactgcgcggccttcggccttgcccaaggtggcgctgccccttggaacccccgcactcgccgccgtgaggctcgggggcaggcgggcgggctt
cgcccttcgactgcccccactcgcataggcttgggtcgttccaggcgcgtcaaggccaagccgctgcgcggtcgctgcgcgagccttgacccgccttccact
tggtgtccaaccggcaagcgaagcgcgcaggccgcaggccggaggcttttccccagagaaaattaaaaaaattgatggggcaaggccgcaggccgcgca
gttggagccggtgggtatgtggtcgaaggctgggtagccggtgggcaatccctgtggtcaagctcgtgggcaggcgcagcctgtccatcagcttgtccagc
agggttgtccacgggccgagcgaagcgagccagccggtggccgctcgcggccatcgtccacatatccacgggctggcaagggagcgcagcgaccgcg
cagggcgaagcccggagagcattgcgcgaaaaggtgagaaaagccgggcactgcccggctttatttttgctgctgcgcgttccaggccgcccacactcgttt
gacctggctcgggctgcatccgaccagcttggccgtcttggcaatgctcgatccgccggagcgaagcgtgatgatgcggtcgtgcatgccggcgtcacgttt
gcggccggtgtagcggccggcggccttcgccaactggacaccctgacgttgacgctcgcgccgatcctcgtagtcgtcgcgggccatctgcaaggcgagc
ttcaaaagcatgtcctggacggattccagaacgattttcgccactccgttcgcctcggcggccagctccgacaggtccaccacgccaggcacggccagcttg
gcccctttggcccggatcgacgcaaccaggcgctcggcctcggccaacggcaagcggctgatgcggtcgatcttctccgcaacgacgacttcaccaggttg
caggtccgcgatcatgcgcagcagctcgggccggtcggcgcgtgcgccggacgccttctcgcggtagatgccggcgacgtagtacccggcggcccgcgt
ggccgctacaaggctctcctggcgttcaagattctgctcgtccgtactggcgcgcaggtagatgcgggcgaccttcaaccttcgtccctccggttgttgctctcg
cgtcgccatttccacggctcgacggcgtgcggatcggaccagaggccgacgcgcttgcctcgcgcctcctgttcgagccgcagcatttcagggtcggccgc
gcggccgtggaagcgataggcccacgccatgccctggtgaaccatcgcggcgttgacgttgcgcggctgcggcggccggctggccagctccatgttgacc
cacacggtgcccagcgtgcggccgtaacggtcggtgtccttctcgtcgaccaggacgtgccggcggaacaccatgccggccagcgcctggcgcgcacgtt
cgccgaaggcttgccgcttttccggcgcgtcaatgtccaccaggcgcacgcgcaccggctgcttgtctaccagcacgtcgatggtgtcgccgtcgatgatgc
gcacgacctcgccgcgcagctcggcccatgccggcgaggcaacgaccaggacggccagcgcggcagcggcgcgcagcatggcgtagcttcggcgctt
catgcgtggccccattgctgatgatcggggtacgccaggtgcagcactgcatcgaaattggccttgcagtagccgtccagcgccacccgcgagccgaacgc
cggcgaaaggtactcgaccaggccgggccggtcgcggacctcgcgccccaggacgtggatgcgccggccgcgtgtgccgtcgggtccaggcacgaag
gccagcgcctcgatgttgaagtcgatggatagaagttgtcggtagtgcttggccgccctcatcgcgtcccccttggtcaaattgggtatacccatttgggcctag
tctagccggcatggcgcattacagcaatacgcaatttaaatgcgcctagcgcattttcccgaccttaatgcgcctcgcgctgtagcctcacgcccacacatgtg
ctaatgtggttacgtgtattttatggaggttatccaatgagccgcctgacaatcgacatgacggaccagcagcaccagagcctgaaagccctggccgccttgc
agggcaagaccattaagcaatacgccctcgaacgtctgttccccggtgacgctgatgccgatcaggcatggcaggaactgaaaaccatgctggggaaccgc
atcaacgatgggcttgccggcaaggtgtccaccaagagcgtcggcgaaattcttgatgaagaactcagcggggatcgcgcttgacggcctacatcctcacg
gctgaggccgaagccgatctacgcggcatcatccgctacacgcgccgggagtggggcgcggcgcaggttcgccgctatatcgctaagctggaacagggc
atagccaggcttgccgccggcgaaggcccgtttaaggacatgagcgaactctttcccgcgctgcggatggcccgctgcgaacaccactacgtttttgcctgc
cgcgtgcgggcgaacccgcgttggtcgtggcgatcctgcatgagcgcatggacctcatgacgcgacttgccgacaggctcaagggctgaccgcgtcagac
gcccgtagcagcccgctacgggctttttcatgcccttgccagcaagcccgtagggcgccgcagccgccgtaggcggtcacgactttgcgaagggccggcc
acctggaaagccacgttgtgtctcaaaatctctgatgttacattgcacaagataaaaatatatcatcatgaacaataaaactgtctgcttacataaacagtaataca
aggggtgttatgagccatattcaacgggaaacgtcttgctcgaggccgcgattaaattccaacatggatgctgatttatatgggtataaatgggctcgcgataat
gtcgggcaatcaggtgcgacaatctatcgattgtatgggaagcccgatgcgccagagttgtttctgaaacatggcaaaggtagcgttgccaatgatgttacaga
tgagatggtcagactaaactggctgacggaatttatgcctcttccgaccatcaagcattttatccgtactcctgatgatgcatggttactcaccactgcgatccccg
ggaaaacagcattccaggtattagaagaatatcctgattcaggtgaaaatattgttgatgcgctggcagtgttcctgcgccggttgcattcgattcctgtttgtaatt
gtccttttaacagcgatcgcgtatttcgtctcgctcaggcgcaatcacgaatgaataacggtttggttgatgcgagtgattttgatgacgagcgtaatggctggcc
tgttgaacaagtctggaaagaaatgcataagcttttgccattctcaccggattcagtcgtcactcatggtgatttctcacttgataaccttatttttgacgaggggaa
attaataggttgtattgatgttggacgagtcggaatcgcagaccgataccaggatcttgccatcctatggaactgcctcggtgagttttctccttcattacagaaac
ggctttttcaaaaatatggtattgataatcctgatatgaataaattgcagtttcatttgatgctcgatgagtttttctaatcagaattggttaattggttgtaacactggca
gagcatt

**Sequence of plasmid pMVRha (8437 bp):**

acgctgacttgacgggacggcgcgccgcttcactcattcgccctagcataaccccttggggcctctaaacgggtcttgaggggttttttgcttaatctttctgcga
attgagatgacgccactggctgggcgtcatcccggtttcccgggtaaacaccaccgaaaaatagttactatcttcaaagccacattcggtcgaaatatcactgat
taacaggcggctatgctggagaagatattgcgcatgacacactctgacctgtcgcagatattgattgatggtcattccagtctgctggcgaaattgctgacgcaa
aacgcgctcactgcacgatgcctcatcacaaaatttatccagcgcaaagggacttttcaggctagccgccagccgggtaatcagcttatccagcaacgtttcgc
tggatgttggcggcaacgaatcactggtgtaacgatggcgattcagcaacatcaccaactgcccgaacagcaactcagccatttcgttagcaaacggcacatg
ctgactactttcatgctcaagctgaccgataacctgccgcgcctgcgccatccccatgctacctaagcgccagtgtggttgccctgcgctggcgttaaatcccg
gaatcgccccctgccagtcaagattcagcttcagacgctccgggcaataaataatattctgcaaaaccagatcgttaacggaagcgtaggagtgtttatcgtca

gcatgaatgtaaaagagatcgccacgggtaatgcgataagggcgatcgttgagtacatgcaggccattaccgcgccagacaatcaccagctcacaaaaatc
atgtgtatgttcagcaaagacatcttgcggataacggtcagccacagcgactgcctgctggtcgctggcaaaaaaatcatctttgagaagttttaactgatgcgc
caccgtggctacctcggccagagaacgaagttgattattcgcaatatggcgtacaaatacgttgagaagattcgcgttattgcagaaagccatcccgtccctgg
cgaatatcacgcggtgaccagttaaactctcggcgaaaaagcgtcgaaaagtggttactgtcgctgaatccacagcgataggcgatgtcagtaacgctggcct
cgctgtggcgtagcagatgtcgggctttcatcagtcgcaggcggttcaggtatcgctgaggcgtcagtcccgtttgctgcttaagctgccgatgtagcgtacgc
agtgaaagagaaaattgatccgccacggcatcccaattcacctcatcggcaaaatggtcctccagccaggccagaagcaagttgagacgtgatgcgctgtttt
ccaggttctcctgcaaactgcttttacgcagcaagagcagtaattgcataaacaagatctcgcgactggcggtcgagggtaaatcattttccccttcctgctgttc
catctgtgcaaccagctgtcgcacctgctgcaatacgctgtggttaacgcgccagtgagacggatactgcccatccagctcttgtggcagcaactgattcagcc
cggcgagaaactgaaatcgatccggcgagcgatacagcacattggtcagacacagattatcggtatgttcatacagatgccgatcatgatcgcgtacgaaac
agaccgtgccaccggtgatggtatagggctgcccattaaacacatgaatacccgtgccatgttcgacaatcacaatttcatgaaaatcatgatgatgttcaggaa
aatccgcctgcgggagccggggttctatcgccacggacgcgttaccagacggaaaaaaatccacactatgtaatacggtcatactggcctcctgatgtcgtca
acacggcgaaatagtaatcacgaggtcaggttcttaccttaaattttcgacggaaaaccacgtaaaaaacgtcgatttttcaagatacagcgtgaattttcaggaa
atgcggtgagcatcacatcaccacaattcagcaaattgtgaacatcatcacgttcatctttccctggttgccaatggcccattttcctgtcagtaacgagaaggtc
gcgaattcaggcgctttttagactggtcgtaatgaacaattcttaagaaggagatatacaaatgtgagacccgcagaaaggcccacccgaaggtgagccagtg
tgactctagtagagagcgttcaccgacaaacaacagataaaacgaaaggcccagtctttcgactgagcctttcgttttatttgaagcttattacacttccagcaca
cgggcaacagcatattcttccagtttgaacgccagaccttctttatccagattcgtgcgggtcagacgcagatccacaaagtggctcggcggcatcagcactg
gtttgttggcgcggtaattgttttcataacgacatttcagcgttttaccgccttcaaccatcaggtacatgatggtgtcgcctttgataataccatcgcacggcgtcat
tttttcagaggtcggttcccagcccagggtgcgtttttgcatcaccggaccatcaatcgggaaattaccgccctgaaagttcgttttatgcaccagggtgttaccct
tcaggctcatttcgaccgtggcggtcgcacaaccgccatcttcaaacgcaattttacgatcgtacgtcaggccttccgggaaggccagtttaaagtagtccgga
atatccgccgggtatttcatgaagcatttgaaaccgtatttcagggtcggagcgacaatatcaaacgcaaacggcagcggaccgcctttcgtaacacgaaagg
tgccactctgtttaccttcatacggtttgccggtaccttcgccttcaatcgtaaaggcgtgaccgttgaccgaaccttccatcagccacgtcattttcatcgtatctg
ccagtgcttgttttgagtgcgacatatgtatatctccttcttaaaagatctaaggtcgatccattaggtttactaacagtatattctaaatttccacctgtgtcaataacg
gtttttatatccgctggtctctgctttttttggcggatgagagaagattttcagcctgatacagattaaatcagaacgcagaagcggtctgataaaacagaatttgcct
ggcggcagtagcgcggtggtcccacctgaccccatgccgaactcagaagtgaaacgccgtagcgccgatggtagtgtggggtgtccccatgcgagagtag
ggaactgccaggcatcaaataaaacgaaaggctcagtcgaaagactgggcctttcgttttatctgttgtttgtcggtgaacgctctcctgagtaggacaaatccg
ccgggagcggatttgaacgttgcgaagcaacggcccggagggtggcgggcaggacgcccgccataaactgccaggcatcaaattaagcagaaggccat
cctgacggatggcctttttttaattaagttccagacaaggtatagggcggcgaggcggctacagccgatagtctggaacagcgcacttacgggttgctgcgcaa
cccaagtgctaccggcgcggcagcgtgacccgtgtcggcggctccaacggctcgccatcgtccagaaaacacggctcatcgggcatcggcaggcgctgc
tgcccgcgccgttcccattcctccgtttcggtcaaggctggcaggtctggttccatgcccggaatgccgggctggctgggcggctcctcgccggggccggtc
ggtagttgctgctcgcccggatacagggtcgggatgcggcgcaggtcgccatgccccaacagcgattcgtcctggtcgtcgtgatcaaccaccacggcggc
actgaacaccgacaggcgcaactggtcgcggggctggccccacgccacgcggtcattgaccacgtaggccgacacggtgccggggccgttgagcttcac
gacggagatccagcgctcggccaccaagtccttgactgcgtattggaccgtccgcaaagaacgtccgatgagcttggaaagtgtcttttggctgaccaccac
ggcgttctggtggcccatctgcgccacgaggtgatgcagcagcattgccgccgtgggtttcctcgcaataagcccggcccacgcctcatgcgctttgcgttcc
gtttgcacccagtgaccgggcttgttcttggcttgaatgccgatttctctggactgcgtggccatgcttatctccatgcggtaggggtgccgcacggttgcggca
ccatgcgcaatcagctgcaacttttcggcagcgcgacaacaattatgcgttgcgtaaaagtggcagtcaattacagattttctttaacctacgcaatgagctattg
cggggggtgccgcaatgagctgttgcgtacccccctttttttaagttgttgatttttaagtctttcgcatttcgccctatatctagttctttggtgcccaaagaagggca
cccctgcggggttcccccacgccttcggcgcggctccccctccggcaaaaagtggcccctccggggcttgttgatcgactgcgcggccttcggccttgccca
aggtggcgctgccccccttggaacccccgcactcgccgccgtgaggctcgggggggcaggcgggcgggcttcgcccttcgactgcccccactcgcataggc
ttgggtcgttccaggcgcgtcaaggccaagccgctgcgcggtcgctgcgcgagccttgacccgccttccacttggtgtccaaccggcaagcgaagcgcgc
aggccgcaggccggaggcttttccccagagaaaattaaaaaaattgatggggcaaggccgcaggccgcgcagttggagccggtgggtatgtggtcgaag
gctgggtagccggtgggcaatccctgtggtcaagctcgtgggcaggcgcagcctgtccatcagcttgtccagcagggttgtccacgggccgagcgaagcg
agccagccggtggccgctcgcggccatcgtccacatatccacgggctggcaagggagcgcagcgaccgcgcagggcgaagtgtagactttccttggtgta
tccaacggcgtcagccgggcaggataggtgaagtaggcccacccgcgagcgggtgttccttcttcactgtcccttattcagagcattgcgcgaaaaggtgag
aaaagccgggcactgcccggctttatttttgctgctgcgcgttccaggccgcccacactcgtttgacctggctcgggctgcatccgaccagcttggccgtcttg
gcaatgctcgatccgccggagcgaagcgtgatgatgcggtcgtgcatgccggcgtcacgtttgcggccggtgtagcggccggcggccttcgccaactgga
caccctgacgttgacgctcgcgccgatcctcgtagtcgtcgcgggccatctgcaaggcgagcttcaaaagcatgtcctggacggattccagaacgattttcgc
cactccgttcgcctcggcggccagctccgacaggtccaccacgccaggcacggccagcttggccccttttggcccggatcgacgcaaccaggcgctcggc
ctcggccaacggcaagcggctgatgcggtcgatcttctccgcaacgacgacttcaccaggttgcaggtccgcgatcatgcgcagcagctcgggccggtcg
gcgcgtgcgccggacgccttctcgcggtagatgccggcgacgtagtacccggcggcccgcgtggccgctacaaggctctcctggcgttcaagattctgctc
gtccgtactggcgcgcaggtagatgcgggcgaccttcaaccttcgtccctccggttgttgctctcgcgtcgccatttccacggctcgacggcgtgcggatcgg
accagaggccgacgcgcttgcctcgcgcctcctgttcgagccgcagcatttcagggtcggccgcgcggccgtggaagcgataggcccacgccatgccctg
gtgaaccatcgcggcgttgacgttgcgcggctgcggcggccggctggccagctccatgttgacccacacggtgcccagcgtgcggccgtaacggtcggtg

tccttctcgtcgaccaggacgtgccggcggaacaccatgccggccagcgcctggcgcgcacgttcgccgaaggcttgccgcttttccggcgcgtcaatgtc
caccaggcgcacgcgcaccggctgcttgtctaccagcacgtcgatggtgtcgccgtcgatgatgcgcacgacctcgccgcgcagctcggcccatgccggc
gaggcaacgaccaggacggccagcgcggcagcggcgcgcagcatggcgtagcttcggcgcttcatgcgtggccccattgctgatgatcggggtacgcca
ggtgcagcactgcatcgaaattggccttgcagtagccgtccagcgccacccgcgagccgaacgccggcgaaaggtactcgaccaggccgggccggtcgc
ggacctcgcgccccaggacgtggatgcgccggccgcgtgtgccgtcgggtccaggcacgaaggccagcgcctcgatgttgaagtcgatggatagaagtt
gtcggtagtgcttggccgccctcatcgcgtccccccttggtcaaattgggtatacccatttgggcctagtctagccggcatggcgcattacagcaatacgcaattt
aaatgcgcctagcgcattttcccgaccttaatgcgcctcgcgctgtagcctcacgcccacacatgtgctaatgtggttacgtgtattttatggaggttatccaatga
gccgcctgacaatcgacatgacggaccagcagcaccagagcctgaaagccctggccgccttgcagggcaagaccattaagcaatacgccctcgaacgtct
gttccccggtgacgctgatgccgatcaggcatggcaggaactgaaaaccatgctggggaaccgcatcaacgatgggcttgccggcaaggtgtccaccaag
agcgtcggcgaaattcttgatgaagaactcagcggggatcgcgcttgacggcctacatcctcacggctgaggccgaagccgatctacgcggcatcatccgct
acacgcgccgggagtggggcgcggcgcaggttcgccgctatatcgctaagctggaacagggcatagccaggcttgccgccggcgaaggcccgtttaagg
acatgagcgaactctttcccgcgctgcggatggcccgctgcgaacaccactacgtttttttgcctgccgcgtgcgggcgaacccgcgttggtcgtggcgatcct
gcatgagcgcatggacctcatgacgcgacttgccgacaggctcaagggctgaccgcgtcagacgcccgtagcagcccgctacgggctttttcatgcccttgc
cagcaagcccgtagggcgccgcagccgccgtaggcggtcacgactttgcgaagggccggccacctggaaagccacgttgtgtctcaaaatctctgatgtta
cattgcacaagataaaaatatatcatcatgaacaataaaactgtctgcttacataaacagtaatacaaggggtgttatgagccatattcaacgggaaacgtcttgc
tcgaggccgcgattaaattccaacatggatgctgatttatatgggtataaatgggctcgcgataatgtcgggcaatcaggtgcgacaatctatcgattgtatggg
aagcccgatgcgccagagttgtttctgaaacatggcaaaggtagcgttgccaatgatgttacagatgagatggtcagactaaactggctgacggaatttatgcc
tcttccgaccatcaagcattttatccgtactcctgatgatgcatggttactcaccactgcgatccccgggaaaacagcattccaggtattagaagaatatcctgatt
caggtgaaaatattgttgatgcgctggcagtgttcctgcgccggttgcattcgattcctgtttgtaattgtccttttaacagcgatcgcgtatttcgtctcgctcaggc
gcaatcacgaatgaataacggtttggttgatgcgagtgattttgatgacgagcgtaatggctggcctgttgaacaagtctggaaagaaatgcataagcttttgcc
attctcaccggattcagtcgtcactcatggtgatttctcacttgataaccttatttttgacgaggggaaattaataggttgtattgatgttggacgagtcggaatcgca
gaccgataccaggatcttgccatcctatggaactgcctcggtgagttttctccttcattacagaaacggctttttcaaaaatatggtattgataatcctgatatgaata
aattgcagtttcatttgatgctcgatgagtttttctaatcagaattggttaattggttgtaacactggcagagcatt
